# Plant litter decomposition in global drylands is better predicted by precipitation seasonality and temperature than by aridity

**DOI:** 10.1101/2025.05.28.655532

**Authors:** Ignacio A. Siebenhart, Pedro M. Tognetti, Agustín Sarquis, Lucio Biancari, Carlos L. Ballaré, Amy T. Austin

## Abstract

Understanding the global carbon (C) balance in terrestrial ecosystems is crucial for predicting their current and future roles as C sources or sinks in the context of global change. Drylands, covering nearly 45% of Earth’s land surface, contribute significantly to net primary production (NPP) and influence the interannual variability of the terrestrial C sink. However, the controls on plant litter decomposition, a major pathway of C release, remain unclear in these ecosystems. Here, we present a global analysis of plant litter decomposition in drylands, using a dataset from 116 sites across five continents spanning diverse climates and ecosystems. We found that litter decomposition does not correlate with mean annual precipitation (MAP) at the global scale, challenging the paradigm that water availability is the primary constraint on ecological processes in drylands. Instead, our analysis identifies mean annual temperature (MAT), precipitation-temperature synchrony, precipitation variability, and cloud cover frequency as key drivers. Specifically, our model predicted faster decomposition rates for warmer and more monsoonal ecosystems, but vary independently of MAP. Additionally, decomposition correlated positively with both lignin and nitrogen content, in contrast to the negative lignin-decomposition relationship commonly observed in mesic ecosystems. These findings suggest a fundamental mismatch between aridity and its expected effects on decomposition rates in terrestrial ecosystems. Given the ongoing expansion of drylands, rising temperatures and changes in precipitation variability under climate change; our results underscore the need to refine decomposition models beyond traditional aridity frameworks. Such refinement is essential for accurately predicting dryland contributions to the global C balance.

## Introduction

The carbon (C) balance in terrestrial ecosystems is primarily determined by inputs from net primary productivity and C outputs to the atmosphere through organic matter decomposition (Schlesinger & Bernhardt, 2020). The C cycle is continuously altered by human activities, primarily through fossil fuel emissions into the atmosphere and land-use changes in terrestrial ecosystems (Calvin et al., 2023). Moreover, feedbacks between the C cycle, climate and human activities complicate our ability to predict the future of C balance in terrestrial ecosystems (Bonan & Doney, 2018; Houghton, 2007). There has been significant progress in understanding the factors regulating C inputs to ecosystems, particularly regarding primary productivity and human-induced modifications of these inputs (Beer et al., 2010; Knapp et al., 2017). In contrast, our understanding of the controls on C release through decomposition remains limited. Plant litter and soil organic matter decomposition are the principal C outputs to the atmosphere, releasing approximately 60 Pg of C as CO_2_ annually; this flux is nearly one order of magnitude larger than the C emissions resulting from human activities (Houghton, 2007; Schlesinger & Bernhardt, 2020). However, decomposition studies have primarily focused on mesic and forest ecosystems (Aerts, 1997; Gholz et al., 2000; Meentemeyer, 1978; Swift et al., 1979; Zhang et al., 2008), leaving a gap in our understanding of this process across terrestrial ecosystems.

Drylands cover ≈45% of the Earth’s land surface (Cherlet et al., 2018; Prăvălie, 2016) and are defined as water-limited ecosystems, where water availability is considered the principal driver of ecological processes (Noy-Meir, 1973; Reynolds et al., 2004). These regions are typically classified as regions with an aridity index (AI), defined as the ratio of mean annual precipitation (MAP) to potential evapotranspiration (PET), below 0.65, indicating that the atmospheric water demand considerably exceeds water supply (Cherlet et al., 2018). As a result, aridity closely aligns with primary productivity, mainly controlled by precipitation amount and variability (Sala et al., 1988; Wang et al., 2022). For carbon release, by contrast, factors such as episodic water pulses and climate variability (Austin et al., 2004; Reynolds et al., 2004), spatial heterogeneity of the vegetation which creates patchy distribution of soil cover and nutrient concentrations (Aguiar & Sala, 1999; Eldridge et al., 2024; Schlesinger & Pilmanis, 1998) and abiotic factors such as photodegradation (Austin & Vivanco, 2006; Rey, 2015) all point to a complex interplay of biotic and abiotic controls on biogeochemical processes.

Drylands play a significant role in the global terrestrial C balance, accounting for approximately 40% of global net primary production (NPP) (Wang et al., 2022) and 15.5% of the total soil organic C (Lal, 2004). Moreover, drylands influence interannual variability and long-term trends in the terrestrial C sink (Ahlström et al., 2015), likely due to the high sensitivity of their NPP to interannual fluctuations in precipitation and temperature (Sasaki et al., 2023). However, discrepancies have been observed between estimated rates of CO_2_ uptake from drylands and the size of organic and inorganic C pools (Schlesinger et al., 2009). These discrepancies could be explained by poorly constrained estimates of C loss, possibly due to unique loss pathways (Austin, 2011; Schlesinger et al., 2009). These complexities highlight the need for accurate assessments of their role as C sinks or sources. Given their widespread distribution and projected expansion due to climate change (J. Huang et al., 2016), a better understanding of C turnover in drylands is essential for predicting their responses to global changes.

Traditional models of plant litter decomposition propose a hierarchical set of controls, with climate and litter quality as primary global-scale controls and the decomposer community dominating at the local-scale (Aerts, 1997; Swift et al., 1979). Climatic factors of temperature and precipitation have traditionally been identified as important, and both positively correlate with decomposition rates across terrestrial ecosystems (Austin & Vitousek, 2000; Gholz et al., 2000; Meentemeyer, 1978). In terms of litter quality, nitrogen (N) content has been positively associated with decomposition, while higher carbon-to-nitrogen (C:N) ratio, lignin content, and lignin-to-nitrogen ratio tend to be associated with lower decomposition rates (Cornwell et al., 2008; Melillo et al., 1982). Recently, this hierarchical approach has been challenged, with new evidence suggesting that the dominance of climate is less important than previously thought. A current view integrates direct and indirect effects of climate affecting decomposition at all scales, from local-scale variability of the decomposer communities (Bradford et al., 2016) to the effects on plant communities and their associated litter quality (Joly et al., 2023).

Most studies on plant litter decomposition have been conducted in forests and mesic ecosystems (Aerts, 1997; Bradford et al., 2016; Joly et al., 2023; Meentemeyer, 1978; Zhang et al., 2008), whereas drylands remain comparatively understudied (Austin, 2011). Several lines of evidence suggest that the controls on litter decomposition in drylands are distinct from those in other terrestrial ecosystems (Austin, 2011; Austin & Vivanco, 2006; Schaefer et al., 1985). In particular, the lack of correlation between litter decomposition and seasonal or mean annual precipitation (Austin, 2011; Vanderbilt et al., 2008), and the fact that traditional models based on climate and litter quality tend to underestimate decomposition rates (Austin, 2011) suggest a complex interplay of controls that are distinct from mesic environments. This complexity likely emerges from the unique combination of conditions of dryland ecosystems, which includes low rainfall, intense solar radiation, pronounced variability in both seasonal and annual precipitation, temperature seasonality, and limited biological activity (Maestre et al., 2016; Whitford & Duval, 2020). Growing evidence suggests that unappreciated controls on C turnover in drylands include exposure to solar radiation, which generates the photochemical breakdown of plant litter (photodegradation) (Austin & Ballaré, 2024; Austin & Vivanco, 2006; Brandt et al., 2010; G. Huang et al., 2017); precipitation-temperature seasonality, which defines alternating periods of biological activity (Davies et al., 2013; Gaxiola & Armesto, 2015); non-rainfall moisture sources (e.g. fog and dew) (Evans et al., 2020); physical fragmentation by macrofauna (Sagi & Hawlena, 2024) and specific interactions with litter quality (Schaefer et al., 1985). Although these controls have been documented at local scales, a global synthesis of climatic drivers for dryland ecosystems at the annual scale is lacking (Cornwell et al., 2008; Zhang et al., 2008). As a result, large-scale patterns and drivers of litter decomposition in drylands remain poorly understood, underscoring the need for a more comprehensive synthesis.

Our objective was to explore climatic and litter quality variables on global patterns of litter decomposition across drylands. For this, we extracted data from the *aridec* database (Sarquis et al., 2022). The resulting dataset used in this study includes 116 dryland sites across six continents, encompassing a wide spatial and climatic range, with MAP from 9 mm to 950 mm and MAT from 1.33°C to 29.06°C (SI Appendix, Fig. S1). We applied a simple exponential model to estimate decomposition rates (*k*, yr^-1^) to regional and global comparisons of decomposition patterns. To identify drivers of decomposition, we selected seven climatic variables from global datasets (Abatzoglou et al., 2018; Wilson & Jetz, 2016) considered relevant for predicting decomposition in drylands (SI Appendix, Fig. S2; Table S2). Mixed-effects regression models were applied to identify key drivers and account for site-specific variability, enhancing the accuracy of variable effect estimations. We also analyzed the relationship between decomposition and litter quality using a subset of the data. Finally, we used the fitted model to extrapolate decomposition rates across global drylands and to describe large-scale patterns.

## Results and discussion

### General global patterns of litter decomposition in drylands

Decomposition rates (*k*) varied widely across drylands, with a global average of 0.75 yr^−1^. Median *k*-values were comparable between H-Arid (including Hyper-Arid and Arid ecosystems; 0.44 yr^−1^) and Semi-Arid sites (0.42 yr^−1^), but were significantly higher in Dry-Subhumid sites (0.81 yr^−1^) (SI Appendix, Fig. S3, Table S3). Monsoon ecosystems exhibited higher median *k* than Mediterranean (SI Appendix, Table S3, see Methods). Surprisingly, there was no significant correlation between *k* and mean annual precipitation (MAP) globally (Fig. 2A). H-arid and Semi-arid sites confirmed this pattern, while Dry-Subhumid sites exhibited a moderate positive correlation (Fig. 2A). Furthermore, the AI was not corelated with *k* across global drylands (SI Appendix, Fig S3B). This was further supported by the observation of similar ranges of decomposition across AI classes, as well as non-significant differences in median *k*-values between H-Arid and Semi-Arid classes (SI Appendix, Table S3). These results indicate that MAP and aridity are not universally robust predictors for litter decomposition in drylands (Austin, 2011), despite being variables that have historically defined dryland ecosystem classification and functioning (Berdugo et al., 2022; Maestre et al., 2016; Noy-Meir, 1973; Reynolds et al., 2004).

At the same time, litter decomposition was not distinctly linked to ecosystem type within dryland ecosystems. Median *k*-values ranged from 0.30 to 0.72 (Fig. 2B; SI Appendix Table S4) and were similar to those observed for temperate forests (Zhang et al., 2008) but lower than those observed in tropical forest ecosystems (Wieder et al., 2009). Moreover, the large range in median *k*-values within each ecosystem type meant that there were comparable decomposition rates across highly different dryland ecosystems (Fig. 2B, SI Appendix Table S4). Notably, decomposition rates observed in desert ecosystems, which were predominantly composed of H-Arid ecosystems, were found to be comparable to those in predominantly Semi-Arid ecosystems such as grasslands and savannas. Furthermore, these rates were significantly higher than those found in forests, which were primarily composed of Semi-Arid and Dry-Subhumid ecosystems (Fig. 2B; Appendix Table S4). Taken together, these results challenge the traditional assumption that higher moisture availability correlates with faster decomposition and suggest that the importance of MAP as a determinant of ecosystem functioning in drylands may be less than what has been previously assumed.

### Climatic variables related to plant litter decomposition across global drylands

We conducted a mixed regression analysis to evaluate the relationship between decomposition rate and seven climate variables (see Methods and SI Appendix, Fig. S2 and Table S2). Using the AIC, we selected the best fitting model, which identified four key variables positively associated with decomposition rates in drylands: mean annual temperature (MAT), temperature-precipitation synchrony (SEASON), precipitation variability (CVp), and frequency of cloudy days (CLOUDC) (Fig. 2, SI Appendix, Table S5). The global model explained an overall 17% of the variance in decomposition (marginal R^2^; SI Appendix, Table S6), with MAT emerging as the strongest predictor, followed by SEASON, CVp, and CLOUDC (Fig. 2; SI Appendix, Table S6). In particular, the model predicted faster decomposition rates in drylands with higher temperatures, greater synchrony between wet and warm seasons (e.g., monsoon ecosystems), higher precipitation variability, and a greater frequency of cloudy days.

To further explore how climatic controls on decomposition vary across dryland types, we conducted additional mixed regression analyses for each aridity class (H-Arid, Semi-Arid, and Dry-Subhumid), as this classification represents a key aspect for understanding and estimating differences in dryland functioning (Berdugo et al., 2020; Maestre et al., 2021). Subset models consistently confirmed the temperature-decomposition relationship observed in the global model, identifying MAT as the most important variable in both H-Arid and Semi-Arid models. In the Dry-Subhumid model, total radiation (RAD) was selected, which may suggest a temperature-related pattern due to high correlation between MAT and RAD (SI Appendix Fig S2). Moreover, SEASON, CLOUDC, and CVp showed varying correlations across aridity classes, suggesting that their contributions to decomposition rates might depend on aridity. SEASON was only important in the H-Arid model, while CLOUDC and CVp were included in the Semi-Arid model but had non-significant relationships (Fig. 2; SI Appendix, Table S7). The H-Arid subset model suggested a non-significant negative relationship between MAP and decomposition. In line with this, the Semi-Arid model identified NDVI, a predictor of vegetation cover that is strongly correlated with MAP, as having a significant negative relationship with decomposition (Fig. 2; SI Appendix, Table S7). These findings provide further evidence of the lack of correlation observed in Fig. 1 and support the exclusion of MAP from the global model.

**Figure 1.**
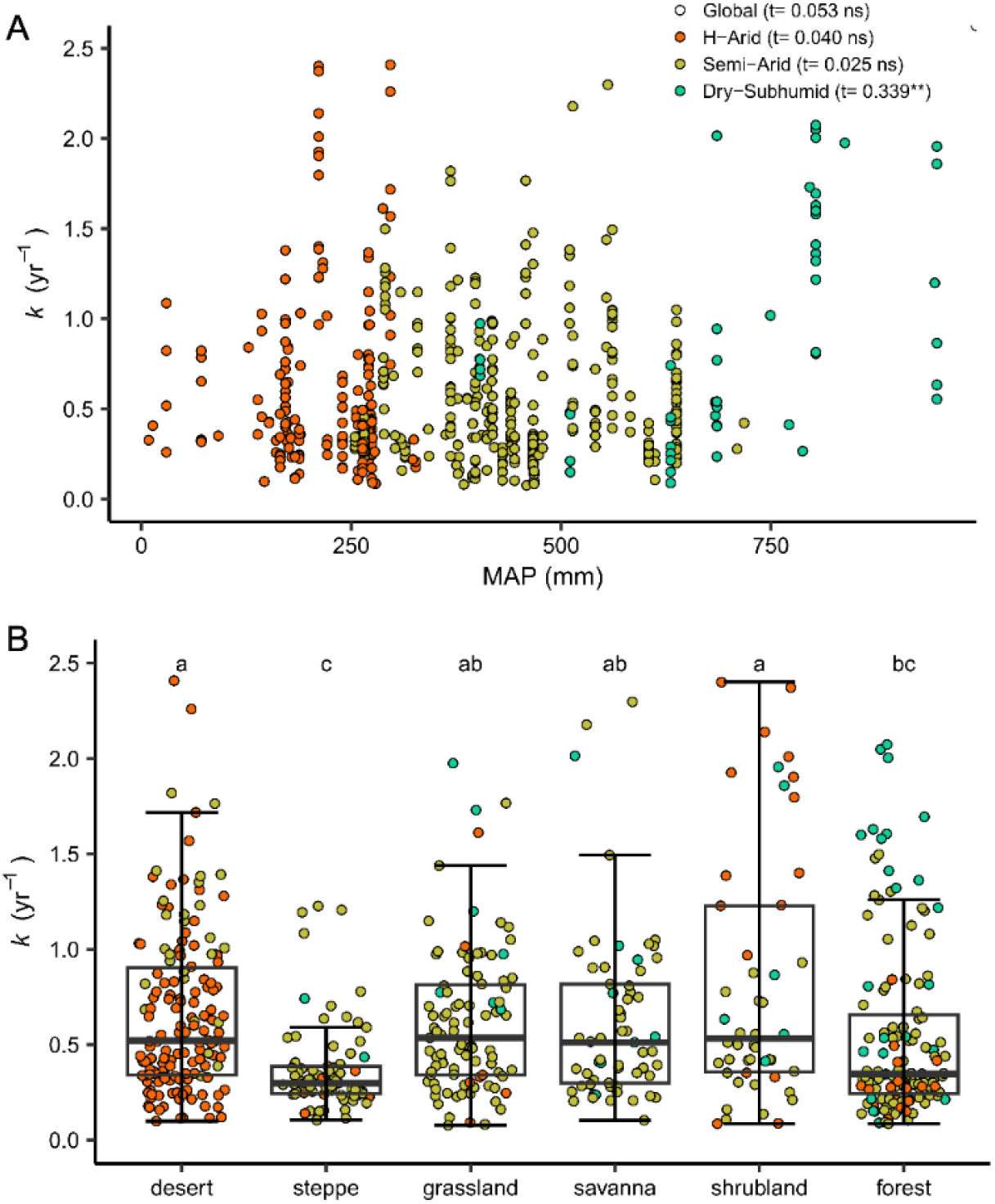
Global patterns of plant litter decomposition in drylands. (A) Annual litter decomposition rate (*k*, yr^−1^) is not correlated with mean annual precipitation (MAP, mm) globally across all aridity classes (t = 0.053), or for hyper-arid (H-Arid) and semi-arid (Semi-Arid) drylands (−0.040, 0.025 ns; respectively), while a moderate positive correlation is observed for the most mesic (Dry-Subhumid) dryland sites (t = 0.339**). Non-parametric Kendall’s correlations are presented for global data and each aridity class (refer to Materials and Methods for details). Significance levels are denoted as **p < 0.01. (B) Decomposition is not strongly linked to ecosystem type across drylands. Ecosystem classifications are based on aridec data (52). Different letters show significant differences in median k-values for different ecosystems from Kruskal–Wallis test corrected by Bonferroni test (refer to Materials and Methods for details, SI Appendix Table S3). Colored symbols in both figures correspond to sites classified by the aridity index (AI) (82) with H-Arid (red); Semi-Arid (yellow-green) and Dry-Subhumid (blue-green).

**Figure 2.**
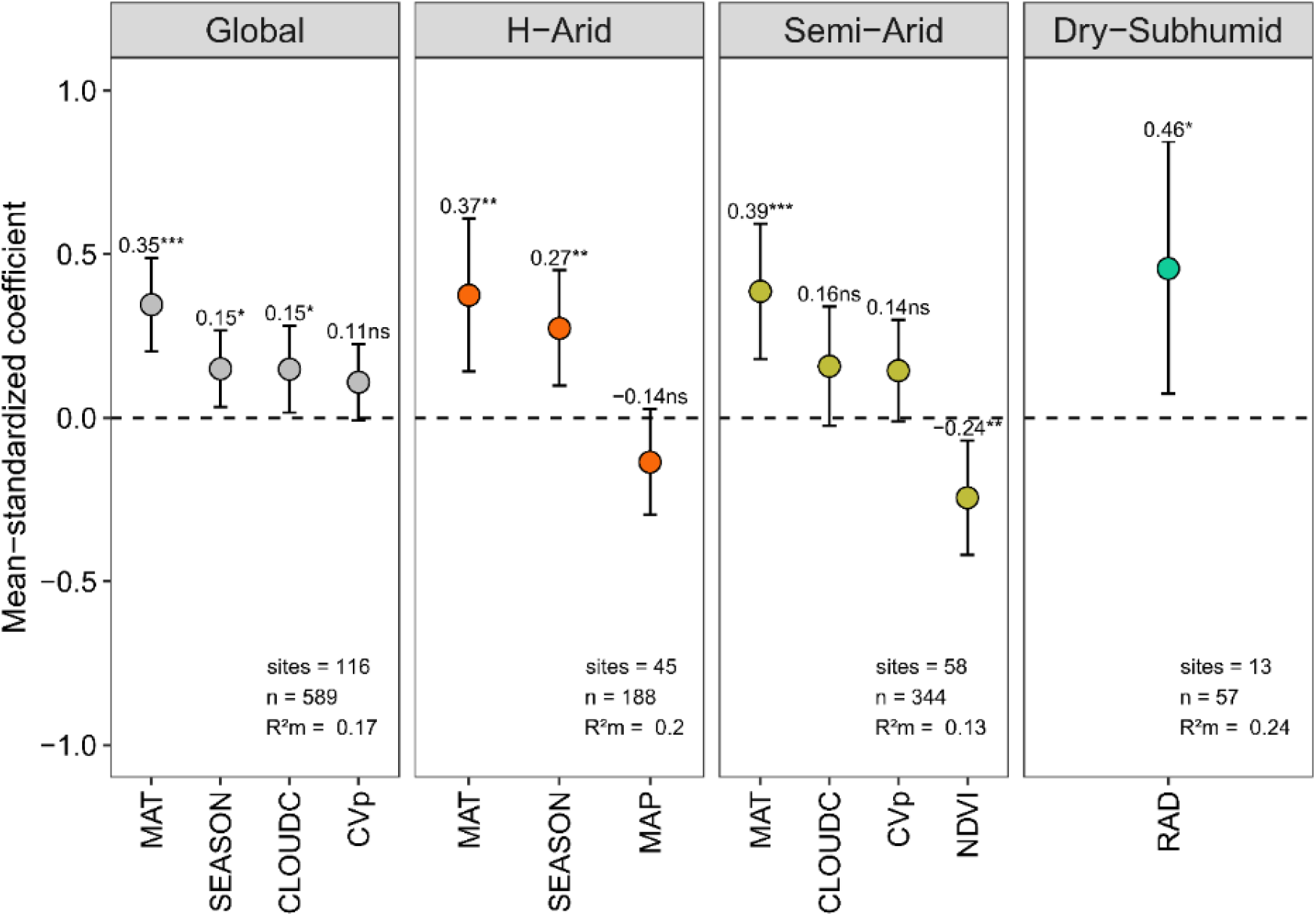
Global and aridity class models demonstrate the importance of temperature, temperature-precipitation synchrony and rainfall variability as predictor variables for litter decomposition in drylands. MAP is not a significant predictor in any model. Globally, mean annual temperature (MAT) is the strongest predictor of litter decomposition, followed by temperature-precipitation synchrony (SEASON) and cloud cover (CLOUDC), all positively correlated with decomposition. Other variables show differential levels of importance in the models depending on the aridity class. Each panel displays the mean-standardized coefficients and their 95% confidence interval (CI) for variables selected in the best linear mixed models for global data and three subsets specific to aridity classification (H-Arid, Semi-Arid, and Dry-Subhumid). Marginal R^2^ (R^2^m) values indicate the percentage of variability explained by fixed effects in each model. Details are provided on the number of sites and observations (n) considered in the analysis (refer to Materials and Methods for details, SI Appendix Table S3-6). Significance levels are denoted as *p < 0.05, **p < 0.01, ***p < 0.001. CVp = precipitation variability, NDVI = mean normalized difference of vegetation index, RAD = annual accumulated radiation.

### Global patterns of plant litter decomposition associated with litter quality in drylands

As we could not include litter quality variables in the general climatic model due to data limitations (see Material and Methods), we instead performed an independent analysis on data subsets where lignin and nitrogen content were available. Decomposition rates showed a positive correlation with both lignin (%) and nitrogen (N) (%) across global drylands (Fig. 2A, B, SI Appendix, Fig S4). Also, lignin:N ratio, a common predictor for decomposition, showed no correlation with decomposition rates for global data and for H-Arid and Semi-Arid drylands (SI Appendix, Fig. S4C). Particularly, lignin content exhibited a threshold effect: decomposition rates increased with lignin content up to 18.5%, but decreased beyond this point, as revealed by a segmented regression (Fig. 3A, SI Appendix Table S8). Correlation analyses also showed a positive relationship between lignin and decomposition, which remained significant within both H-Arid and Semi-Arid classes (SI Appendix Fig. S4A). Additionally, after accounting for the variability explained by climatic factors, lignin showed no significant correlation with the residuals of the global climatic model (SI Appendix Fig. S5A). In the case of N content, it showed a consistent positive relationship with decomposition rates across global drylands (Fig. 3B, SI Appendix Fig. S4B, Table S9), and the correlation remained significant across all aridity classes, although weaker in more arid environments. This positive relationship persisted even after adjusting for climatic variability (SI Appendix Fig. S5B).

**Figure 3.**
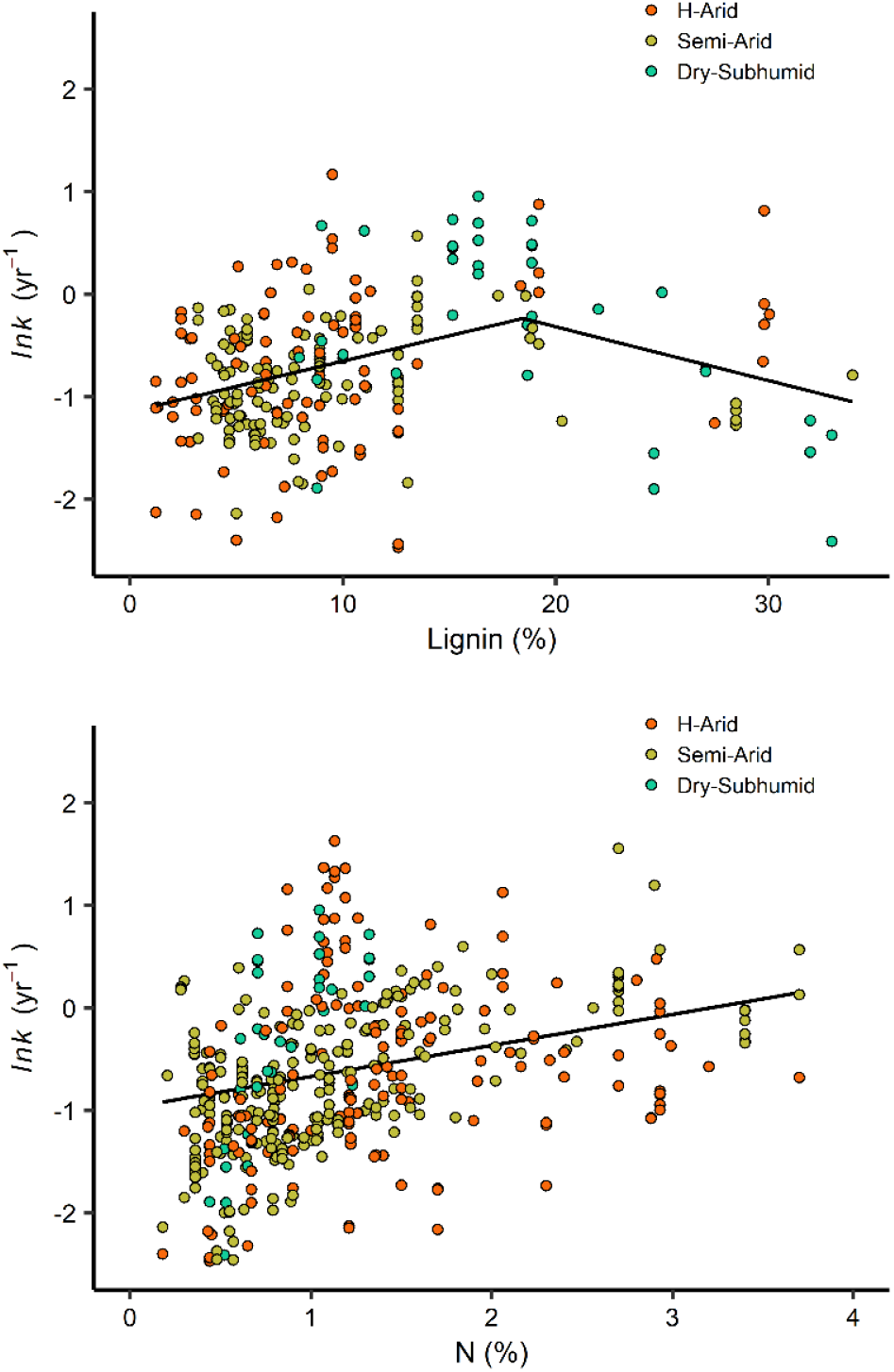
Litter decomposition is positively associated with lignin (A) and nitrogen (N) content (B) globally in drylands. (A) Segmented regression of initial lignin content of litter with decomposition across global drylands shows significant and positive correlation for low values of lignin (<18.5%) but a negative relationship at higher lignin content. (B) Initial N content correlates positively and linearly with litter decomposition across global drylands. Mixed linear models were performed. Litter decomposition rates (*k*) were log-transformed before analysis. Details of statistical analysis can be found in Material and Methods and SI, Appendix, Table S7-8. Color codes for AI classes follow Fig. 1.

### Geographical global patterns of litter decomposition in drylands worldwide

To describe the global geographic patterns of litter decomposition in drylands, we estimated *k* rates globally by extrapolating from our best-fitting model, using climatic layers of the four selected variables. The resulting patterns revealed a strong latitudinal trend, with higher decomposition rates in tropical and subtropical areas and lower rates at higher latitudes, consistent with the importance of MAT as a significant variable in the global model (Fig. 4A). Additionally, monsoon ecosystems exhibited higher decomposition rates than Mediterranean ecosystems, with noticeable changes in *k* occurring even with minor latitudinal shifts (Fig. 4A, SI Appendix Fig. 6). The variables CLOUDC and CVp appeared to fine-tune decomposition rates within the same latitudinal range (Fig. 4A, SI Appendix Fig. 6). To further assess these patterns, we conducted a random sampling of 12,000 points across drylands (SI Appendix Fig. 7). This analysis confirmed that predicted decomposition rates increased consistently with temperature and precipitation-temperature synchrony (SEASON), particularly in warmer drylands, where monsoon ecosystems showed higher values compared to Mediterranean ones (Fig. 4D).

**Figure 4.**
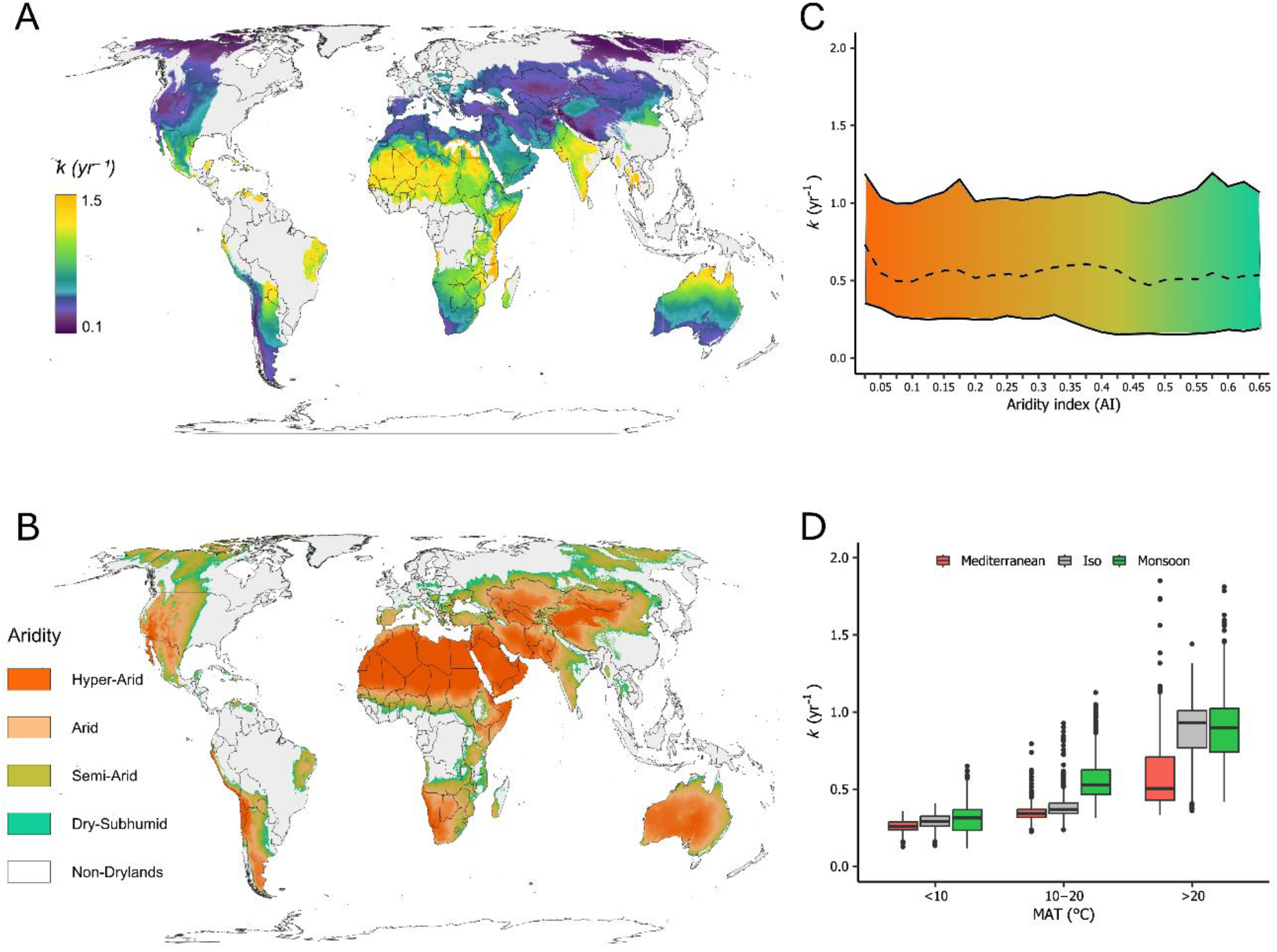
Annual plant litter decomposition across global drylands do not coincide with aridity. (A) Geographical patterns of decomposition rate (*k*; yr^−1^) are estimated using the empirical model developed in this study. (B) Global distribution of drylands based on (Trabucco & Zomer, 2018) data, with AI classes. Comparison between maps A and B demonstrates little coherence between rates of decomposition and aridity classes. (C) *k* values across all AI classes as predicted by our model for global drylands. The solid lines represent the 5th and 95th percentiles of *k* for predicted data grouped into AI 0.025 intervals, and the dashed line represents the mean value of *k*. Annual decomposition rates show similar predicted values across the aridity range. (D) Alternative grouping of litter decomposition by MAT and SEASON demonstrates consistent patterns of increased decomposition with temperature and in monsoon-type ecosystems. Decomposition values from both (C) and (D) are derived from a random sampling of 12,000 predicted values by our global model in global drylands (refer to Materials and Methods for details, SI Appendix, Fig. S5). Color scales for (A-D) are shown in each panel.

Geographical patterns clearly showed the mismatch observed between decomposition and aridity where litter decomposition across global drylands did not coincide with aridity classes (Fig. 4A-B). Our model predicted similar decomposition rates across the entire aridity range (Fig. 4C), suggesting that aridity alone was not a strong determinant of litter decomposition. Instead, decomposition rates were highest in regions where warm temperatures coincided with lower aridity, particularly in semi-arid and dry-subhumid subtropical zones. Conversely, in areas where lower aridity did not coincide with higher temperatures, decomposition rates were low (Fig. 4B, SI Appendix Fig. 6).

### Toward a broader understanding of C turnover in dryland ecosystems

Understanding the mechanisms driving C cycling in drylands is essential for improving C flux models, evaluating the dynamics of soil organic matter, and predicting decomposition and long-term C storage. Our global models identified MAT as the most important climate variable across drylands as well as within different aridity classes (Fig. 2). In line with previous empirical evidence of the importance of temperature on litter decomposition (Gholz et al., 2000; Liski et al., 2003; Zhang et al., 2008), as well as recent global syntheses (Djukic et al., 2018; Wu et al., 2025; Zhao et al., 2025), our model projected patterns of plant litter decomposition that were strongly associated with latitudinal changes in temperature (Fig. 4A). Nevertheless, our global model detours substantially from other models (Djukic et al., 2018; Wu et al., 2025; Zhao et al., 2025), identifying new insights into the linkages between climatic predictors and ecosystem functioning as well as litter quality relationships in dryland ecosystems.

Perhaps most noteworthy in this analysis was that MAP did not correlate with annual rates of litter decomposition (Fig. 1), nor was it included as a significant variable in any of the models by aridity class (Fig. 2). This lack of correlation between MAP and decomposition rates confirms empirical studies at local or regional scales in drylands (Austin, 2011; Vanderbilt et al., 2008) at a global scale, but contrasts with predictive models of climate-driven decomposition from empirical studies (Austin & Vitousek, 2000; Gholz et al., 2000; Meentemeyer, 1978) as well as global analyses (Wu et al., 2025; Zhang et al., 2008; Zhao et al., 2025).

Why is MAP a poor predictor of plant litter decomposition in drylands? Our results highlight underappreciated aspects of rainfall in drylands, including seasonality and interactions with temperature that may explain mismatches with traditional models of drylands functioning. In addition, the constraints from water scarcity on biotic activity may increase the importance of abiotic drivers of decomposition such as photodegradation (Austin, 2011; Rey, 2015). Drylands are characterized by high rainfall variability and seasonality, with precipitation concentrated in specific periods (Noy-Meir, 1973; Wang et al., 2022; Whitford & Duval, 2020), often more markedly than in other ecosystems (Maestre et al., 2021; Whitford & Duval, 2020). These wetter seasons coincide with increased biotic activity and higher decomposition rates (Berenstecher et al., 2020; Davies et al., 2013; Gaxiola & Armesto, 2015; Hewins & Throop, 2016), consistent with the marginally positive effect of intra-annual variability we observed (Fig. 2). This also aligns with findings that precipitation at a monthly scale correlates with short-term decomposition (Zhao et al., 2025), in contrast to the lack of correlation at the annual scale. Moreover, synchrony between warm and wet seasons also emerged as a key driver: drylands with high coincidence showed higher decomposition rates than Mediterranean climates, where rainfall occurs in cooler periods (Fig. 2, 4A, D). This temporal mismatch in Mediterranean systems leads to seasonal shifts in dominant controls: biotic decomposition in winter and abiotic in summer (Berenstecher et al., 2020). Whereas monsoonal drylands show intense biotic pulses during humid summers (Hewins & Throop, 2016). These patterns were particularly relevant in very arid systems with high maximum temperatures (Fig. 4D; SI Appendix Fig. S5). Taken together, our findings suggest that timing and variability of precipitation play a much more important role than MAP in driving decomposition dynamics in global drylands.

The potential for confounding covariation can be important in the interpretation of the global models presented here and elsewhere (Bradford et al., 2016; Wu et al., 2025; Zhao et al., 2025). For instance, the covariation between MAT and other climatic variables may obscure the influence of those variables on litter decomposition. While MAP and MAT showed a weak correlation across the dataset, MAT was strongly correlated with annual accumulated radiation (RAD) (Appendix S, Figs. S1, S2). For example, the strong co-variation of solar radiation and MAT complicates the separation of their individual effects –for example, the potential importance of photodegradation increasing decomposition in hot zones with very high solar irradiance. Given that photodegradation is a recognized control of decomposition in drylands (Austin & Ballaré, 2024; Austin & Vivanco, 2006; Brandt et al., 2010; Day et al., 2007), part of the observed positive relationship with higher temperatures could be linked to increased solar irradiance. Furthermore, as lignin is positively and negatively associated with litter decomposition, retarding biotic decomposition but promoting photodegradation (Austin et al., 2016; Austin & Ballare, 2010), the positive correlation between decomposition rates and lignin content (Fig. 3A, SI Appendix Fig.

S4A) supports the hypothesis that part of the positive correlation between *k* and MAT may be caused by increased solar radiation and hence photodegradation, rather than by a thermal acceleration of biotic activity. However, estimating photodegradation at a global scale is challenging, as solar radiation from climate databases do not always accurately reflect litter exposure to sunlight (Austin & Ballaré, 2024; Liu et al., 2015; Méndez et al., 2022). Since incorporating photodegradation improves predictions of litter decomposition in drylands (Adair et al., 2017), better proxies for litter exposure to solar radiation are essential for quantifying its global impact.

Our results highlight both contrasting and convergent patterns in how litter quality influences decomposition rates in global drylands. In contrast to the negative relationship between lignin content and decomposition rates reported for mesic ecosystems (Cornwell et al., 2008; Meentemeyer, 1978; Melillo et al., 1982), our analysis showed a distinct pattern in global drylands (Fig. 2A; SI Appendix, Fig S4A). Positive correlation and the non-linear pattern for lignin confirms previous evidence of unique patterns in these ecosystems (Liu et al., 2015; Schaefer et al., 1985; Vanderbilt et al., 2008) where lignin could play a dual role in decomposition processes (Austin et al., 2016; Austin & Ballare, 2010). The lower biotic activity in drylands (Maestre et al., 2016; Noy-Meir, 1973), combined with the relative importance of photodegradation (Austin & Ballaré, 2024), could explain our results. In contrast, N consistently showed a positive relationship with decomposition which aligns with many previous frameworks derived from mesic ecosystems and global synthesis data. Even under dryland constraints, N remains an important driver, which is consistent with evidence derived from local studies in drylands (Davies et al., 2013; Martínez-Yrízar et al., 2007). However, the relationship may be modulated by distinctive patterns of nitrogen availability in drylands, influenced by seasonal water pulses and nitrogen turnover (Austin et al., 2004; Parton et al., 2007). These patterns likely reflect both direct biotic effects and indirect consequences of other factors that shape litter quality.

Although aridity is widely used to classify and predict functional attributes of dryland ecosystems (Berdugo et al., 2022; Maestre et al., 2016, 2021) our global analysis suggests that it is a poor predictor of the annual rate of litter decomposition. However, to estimate the actual release of C, one must also consider the amount of biomass that could decompose, which is largely determined by NPP and, in turn, closely related to MAP (Knapp et al., 2017; Sala et al., 1988; Wang et al., 2022). Unearthing this key mismatch between aridity and the velocity of C release advances our mechanistic understanding of dryland ecosystem functioning and has important implications for understanding their carbon balance. These results challenge long-standing assumptions about the dominant controls on C cycling in these ecosystems, which are based nearly exclusively on water availability (Maestre et al., 2021; Noy-Meir, 1973; Wang et al., 2022). Considering aridity alone may mislead our predictions in these ecosystems, particularly by underestimating decomposition rates and overestimating C storage (Austin, 2011; Schlesinger et al., 2009). A more comprehensive understanding of these alternative drivers is crucial for improving C cycle models and accurately assessing the role of drylands in the global C balance.

Understanding controls on litter decomposition is important as we move more concretely into an era of realized climate change with alterations in biogeochemical cycle at most temporal and spatial scales (Calvin et al., 2023). Given the importance of drylands in the global C cycle, both due to their vast extent and their influence on interannual C variability (Ahlström et al., 2015; Prăvălie, 2016; Wang et al., 2022), understanding how rising temperatures will impact drylands is necessary for assessing their role in the global C balance. While higher temperatures could enhance biotic decomposition, they are also expected to increase atmospheric evaporative demand (ETP), potentially intensifying aridity (Grünzweig et al., 2022; J. Huang et al., 2016). At the same time, our findings suggest that aridity alone will not be the main driver of changes in litter decomposition. Instead, changes in the seasonality of Mediterranean- and monsoon-type ecosystems and increasing intra-annual rainfall variability may play a more central role in shaping decomposition dynamics. The large and unpredictable shifts in precipitation variability and seasonality projected under climate change (Calvin et al., 2023) and the unappreciated responses of ecosystems to these changes (Hajek & Knapp, 2022) could further challenge our ability to anticipate dryland ecosystem responses. Moreover, as global climate patterns shift, ecosystems once considered mesic may exhibit more dryland-specific mechanisms (Grünzweig et al., 2022), and together with the projected expansion of drylands (J. Huang et al., 2016), this underscores the need for a more comprehensive understanding of decomposition controls in global drylands.

## Materials and Methods

To assess global patterns of plant litter decomposition and aim our objectives, we compiled a comprehensive dataset that includes decomposition data, climatic variables, and litter quality indicators from various global and field-based sources.

### Decomposition data

We extracted litter decomposition data from *aridec*, a global database on litter mass loss from drylands ecosystems (Sarquis et al., 2022). The *aridec* database comprises 1,752 time series of litter mass loss corresponding to 184 publications from 212 dryland sites across six continents, representing a wide range of climates and biomes (Sarquis et al., 2022). Data were considered to be from the same site when tagged with identical geographic coordinates. Due to the inherent heterogeneity in *aridec*’s data, a conservative pre-filtering process was applied to ensure data consistency and relevance. First, we excluded data from sites with an aridity index (AI) above 0.65, as sites above this threshold are classified as humid. Lastly, we excluded mass loss series associated with manipulative treatments (e.g., irrigation, nutrient addition, biocide application) and only included those from litterbags placed at the soil surface. The resultant data included 116 sites corresponding to 105 published studies with 588 mass loss time series (Fig. 1A; SI Appendix, Table S1).

We calculated the decomposition rate (*k*) for each one of the mass loss time series using a simple negative exponential model (Olson, 1963):

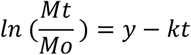

Where Mt is mass remaining at time t, M_0_ is initial mass, k is the decomposition rate. *k* values were calculated with the *nls* function of the R stats package. Because decomposition data included some time series of litter mass loss with an extension of less than a year, we decided to obtain k values per month. For analyses and visualization, data were converted to year.

### Climate data

We selected seven candidate predictor variables from global datasets (SI Appendix, Table S2). Three main variables were obtained from TerraClimate 1981-2010 climatologies (Abatzoglou et al., 2018): mean annual precipitation (MAP), mean annual temperature (MAT), and total solar radiation (RAD). Additionally, we derived two variables representing seasonality from the same dataset: the precipitation-temperature synchrony (SEASON) and annual precipitation variability (CVp). SEASON represents the overlap between wet and warm seasons, which we estimated with the Pearson correlation coefficient on monthly data of precipitation and temperature (Yahdjian et al., 2021). SEASON values range between -1 and 1, where negative values correspond to Mediterranean climates with dry summers and rainy winters, while positive values correspond to Monsoon climates with rainy summers and dry winters. CVp reflects the intra-annual variability of precipitation, based on the coefficient of variation of monthly values, with higher values indicate stronger rainfall seasonality.

Furthermore, we included two predictor variables derived from satellite data products: the Normalized Difference Vegetation Index (NDVI) and CloudCover (CLOUDC). NDVI represents the annual average with higher NDVI values positively correlated with vegetation cover. NDVI data were extracted from The Land Processes Distributed Active Archive Center (LP DAAC) and correspond to the MODIS product (VNP13A3 v001). CloudCover represents the annual mean frequency of days with cloud cover, obtained from EarthEnv (Wilson & Jetz, 2016). A correlation matrix between all variables is presented at SI Appendix, Fig S2. We also obtained the aridity index (AI) as the ratio of MAP to potential evapotranspiration (PET) from CGIAR database (Trabucco & Zomer, 2018), with higher AI indicating less aridity.

### Litter quality data

Lignin and Nitrogen (N) contents were selected as indicators of litter quality.

These data were primarily sourced from *aridec* (Sarquis et al., 2022) whenever available. In cases where data were incomplete, we completed them with values from the same species sourced from other publications. For lignin, we specifically included only those measurements obtained via the acid digestion method (ADL-Lignin) to account for methodological differences that can affect results (Hatfield & Fukushima, 2005). The resulting dataset comprises distinct subsets for each variable:

Lignin (n=254, sites=60) and N (n=452, sites=86).

### Classification of data and exploration

We classified the data according to aridity, ecosystem type, and seasonality. For the aridity classification, we utilized standard classes based on the aridity index. Due to the limited number of hyper-arid sites (AI < 0.05) in the database, we combined them with arid sites 0.05 > AI < 0.2) into a single class named H-Arid. The classes are as follows: H-Arid, (AI < 0.2); Semi-arid (0.2 ≤ AI < 0.5); and Dry-Subhumid (0.5 ≤ AI < 0.65). H- Arid and Semi-Arid classes, which represent the majority of global drylands, were dominant in our dataset (SI Appendix, Table S2). Ecosystem classification was based on aridec data (Sarquis et al., 2022) and corresponded to the ecosystem types reported by the authors of the studies. These types were simplified into six classes: desert, steppe, grassland, savanna, shrubland, and forest. Reflecting the pronounced climatic seasonality typical of drylands, we divided sites based on SEASON values into Mediterranean (−1 to -0.3), Iso (−0.3 to 0.3), and Monsoon (0.3 to 1) ecosystems. Mediterranean ecosystems are characterized by a lack of synchrony between higher monthly precipitation and temperatures, with precipitation occurring during cooler months. In contrast, Monsoon ecosystems exhibit a high coincidence of warm and rainy seasons. Iso ecosystems display uniform patterns in the precipitation-temperature synchrony.

### Data analyses

All statistical analyses were performed in the R, version 4.4.0 (R Core Team, 2023).

#### Exploratory analyses

We performed a principal component analysis (PCA) to define site distribution across climate space, characterized by seven climate variables. We applied Kendall’s correlation to examine relationships between decomposition and MAP globally and within specific aridity classes. Additionally, to explore differences in decomposition rates across aridity, ecosystems and seasonality classes, we compared decomposition rates between classes using a non-parametric Kruskal-Wallis test with Bonferroni correction.

#### Evaluation of relationship between climate and decomposition rate (k)

To evaluate the influence of climatic variables on the decomposition rates we fitted linear mixed models (lmm) using nlme package (Bates et al., 2015). Climatic variables were considered as fixed effects and sites (n=116) were considered as random factors (Appendix SI, Table S1). All predictors were standardized before analyses. We chose best-fit models considering the Akaike Information Criterion (AIC) using the Mumin package (Bartoń, 2010). Partial R^2^ (Part R^2^) and inclusive R^2^ (IR^2^) were obtained for each explanatory variable using the partR^2^ package (Stoffel et al., 2021). Part R^2^ gives an estimation of the proportion of the variance in the response variable determined by an explanatory variable while accounting for covariance between this variable and the other explanatory variables in the model. IR^2^ gives an estimation of the contribution to linear explanatory variables independent of the others (Stoffel et al., 2021).

#### Evaluation of relationship between litter quality and decomposition rate

Due to limited data availability, we were unable to include litter quality variables in the climatic global analyses. First, we applied Kendall’s correlation to examine relationships between decomposition and Lignin, Nitrogen contents and L:N ratio globally and within specific aridity classes. In addition, to evaluate the specific relationship between litter quality variables and decomposition we fitted simple lmm specifically for nitrogen and for lignin. In the case of lignin, initial exploration of the data revealed a non-linear relationship with decomposition rates, characterized by a marked threshold: decomposition increased with lignin content up to a certain point, after which the relationship became negative. To capture this pattern, we first identified the breakpoint and then applied a segmented regression model, which allows for different slopes below and above the threshold.

This approach was particularly suitable to reflect the dual role of lignin in dryland decomposition. Also, we examined correlations between N, Lignin contents and the residuals of the global climate model. Using Kendall correlation coefficients, we tested the relationship between these residuals and each litter quality variable. This approach enabled us to evaluate the degree of residual variability not explained by the climate model and its potential association with litter quality independent of climate.

### Geographic patterns of plant litter decomposition across global drylands

We estimated decomposition rates for global drylands by extrapolating our global model to each dryland pixel using QGIS version 3.34.1 (QGIS Development Team, 2021). Using climate layers from TerraClimate, we generated layers for the four climate variables relevant to dryland decomposition. These layers were then integrated with the global decomposition model and the distribution of global drylands to create a comprehensive map of litter decomposition patterns worldwide. Additionally, we generated individual maps for each climate variable as well as for aridity. To further illustrate these patterns, we performed a random sampling of predicted global decomposition rates at 12,000 points, along with the climate data associated with each point. This random sampling allowed us to more clearly depict the decomposition patterns predicted by our model in relation to aridity and climate variables.

## Data Availability

Datasets and code from this study are freely available from figshare: https://figshare.com/s/b4ed61945eddb5b2d23c

## Acknowledgments

I.A.S was supported by a doctoral fellowship from Consejo Nacional de Investigaciones Científicas y Técnicas (CONICET), Argentina and received partial funding from the Neotropical Grassland Conservancy (NGC). Support for research came from grants from the Universidad de Buenos Aires (UBACyT), Argentina (A.T.A and C.L.B); the Fondo para la Investigación Científica y Tecnológica (FONCYT; Agencia I+D+I, Argentina) and the New Phytologist Foundation, United Kingdom (A.T.A and C.L.B). Additional support came from Fondation L’Oréal for Women in Science, France (A.T.A) and the Alexander Von Humboldt Stiftung, Germany (A.T.A. and I.A.S). We thank M. Oesterheld and S. Méndez for their valuable comments during the early stages of this manuscript.

