## Supplemental Information for "Plant litter decomposition in global drylands is better predicted by precipitation seasonality and temperature than by aridity"

**This PDF file includes:**

Figures S1 to S7
Tables S1 to S4
SI References

**Fig. S1. Geographical and climate distribution of sites.** (A) Geographical distribution of sites. Points indicate the location of sites included in analyses. Point color corresponds to three categories derived from aridity index (IA) previously defined: H-Arid, Semi-Arid, Subhumid. (B-C) Climate space defined by the seven selected climate variables (see SI Appendix Table 2). Axes represent the first two components (PC1 and PC2) of the principal components analysis for climate variables; sites organized from colder (left) to warmer along PC1 axis and from drier to more humid along PC2; seasonality patterns for climate space are explained by PC3 and PC4. MAT= Mean annual temperature, MAP= mean annual precipitation, RAD= Total solar radiation, CLOUDC = frequency of cloudy days, NDVI= mean annual normalized difference in vegetation index, CVp= precipitation variability estimated as coefficient of variation for precipitation, SEASON= Seasonality defined as the synchrony of monthly temperature and precipitation.

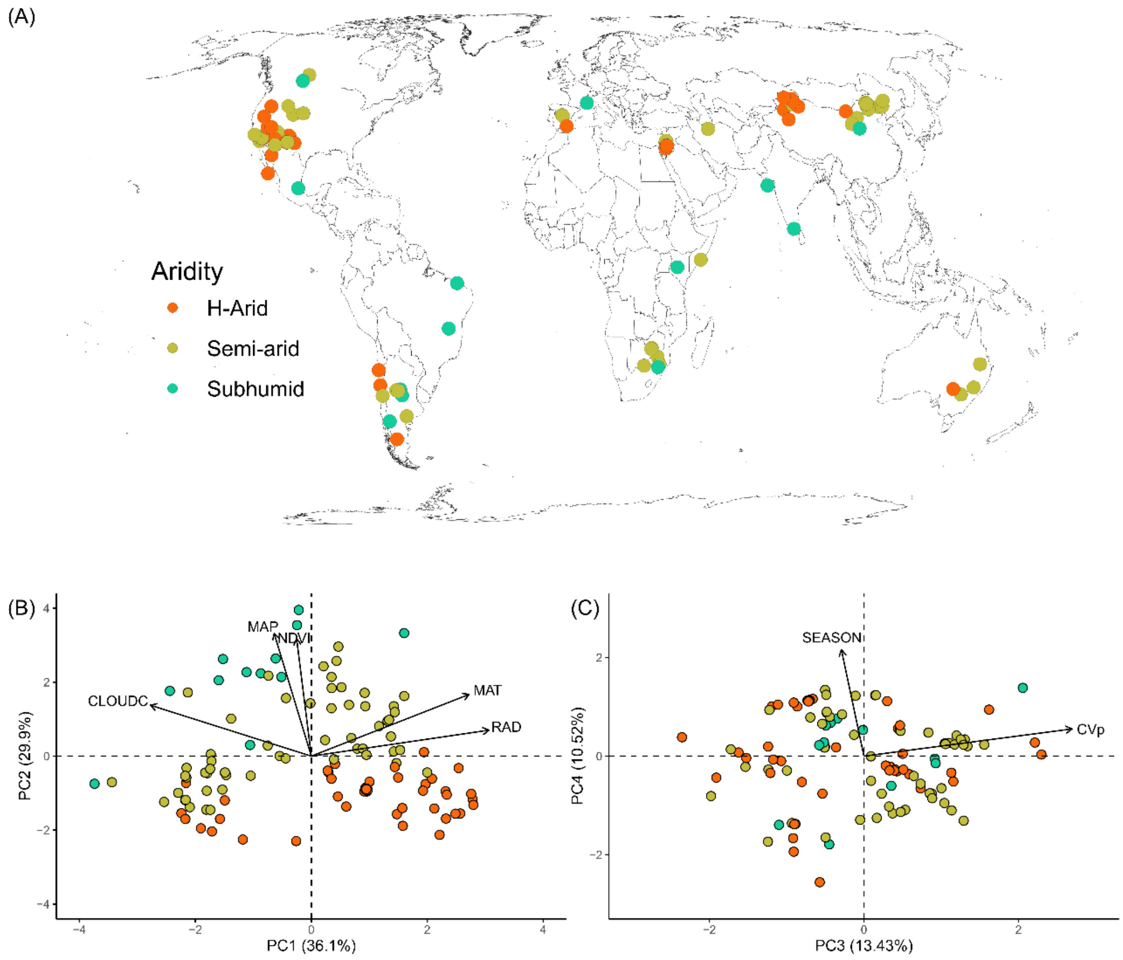

**Fig. S2. Correlation matrix between climate variables.** The diagonal panel shows each variable distribution. The upper panels show spearman Kendall correlation coefficient for each relationship, and the lower panels show pairwise scatter plots between variables. Correlations are significant with p<0.05*; p <0.01**; p<0.001***. MAT= Mean annual temperature, MAP= mean annual precipitation, RAD= Total solar radiation, CLOUDC = frequency of cloudy days, SEASON= Seasonality defined as the synchrony of monthly temperature and precipitation, CVp= precipitation variability estimated as coefficient of variation for precipitation, NDVI= mean annual normalized difference in vegetation index, CLOUDC = frequency of cloudy days.

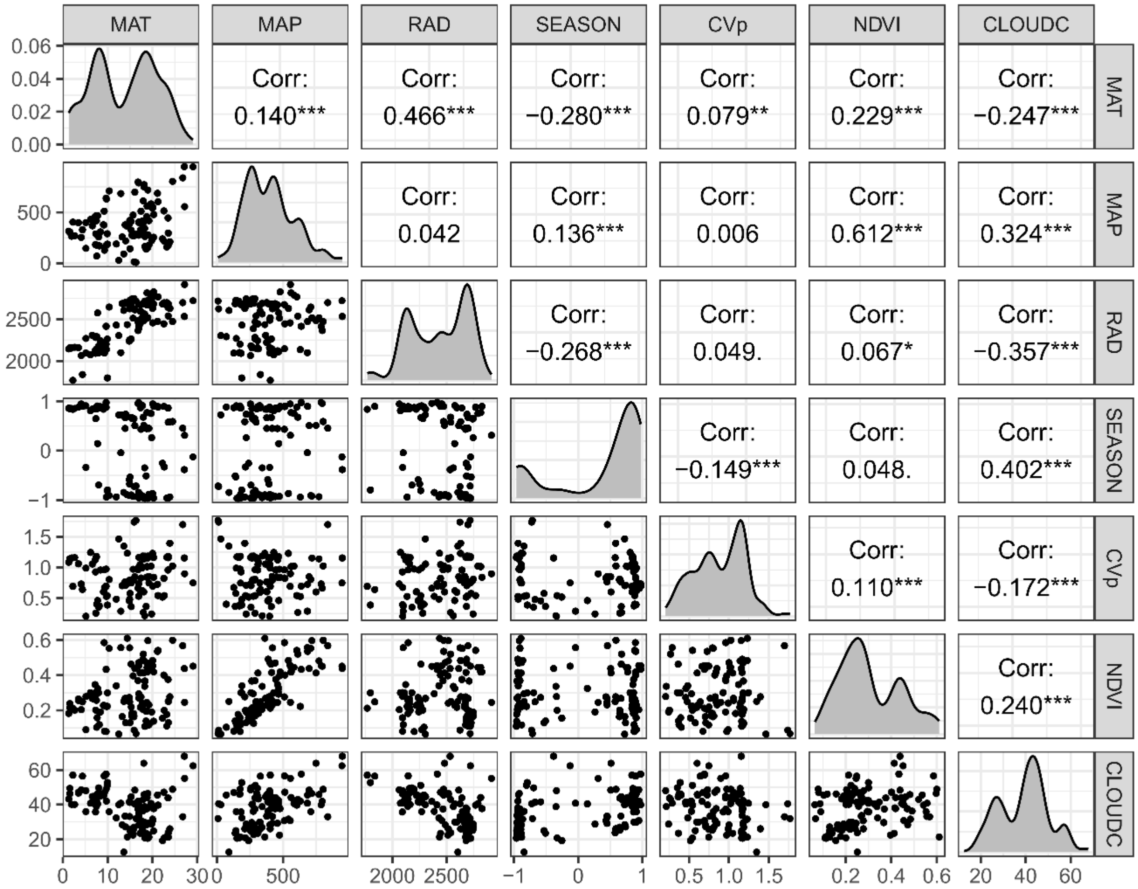

**Fig. S3. Decomposition patterns related to Aridity Index (AI). (A) Decomposition is not strongly related to aridity classes.** H-Arid and Semi-Arid drylands have similar median decomposition rates (*k*). Different letters show significant differences in median *k*-values for different AI classes from Kruskal–Wallis test corrected by Bonferroni test (refer to Materials and Methods for details, SI Appendix Table S3). (B) Relationship between *k* (yr-1) and Aridity Index (AI) for global drylands. Aridity is not correlated with decomposition rates (t=0.023, p-value= 0.420 non-significant). Dashed lines represent superior limit of AI categories. Non-parametric Kendall's correlations are presented for global data and each aridity class (refer to Materials and Methods for details). Colored symbols and lines correspond to sites classified by the aridity index (AI, Trabucco & Zommer, 2018) with H-Arid (red); Semi-Arid (yellow-green) and Dry-Subhumid (blue-green).

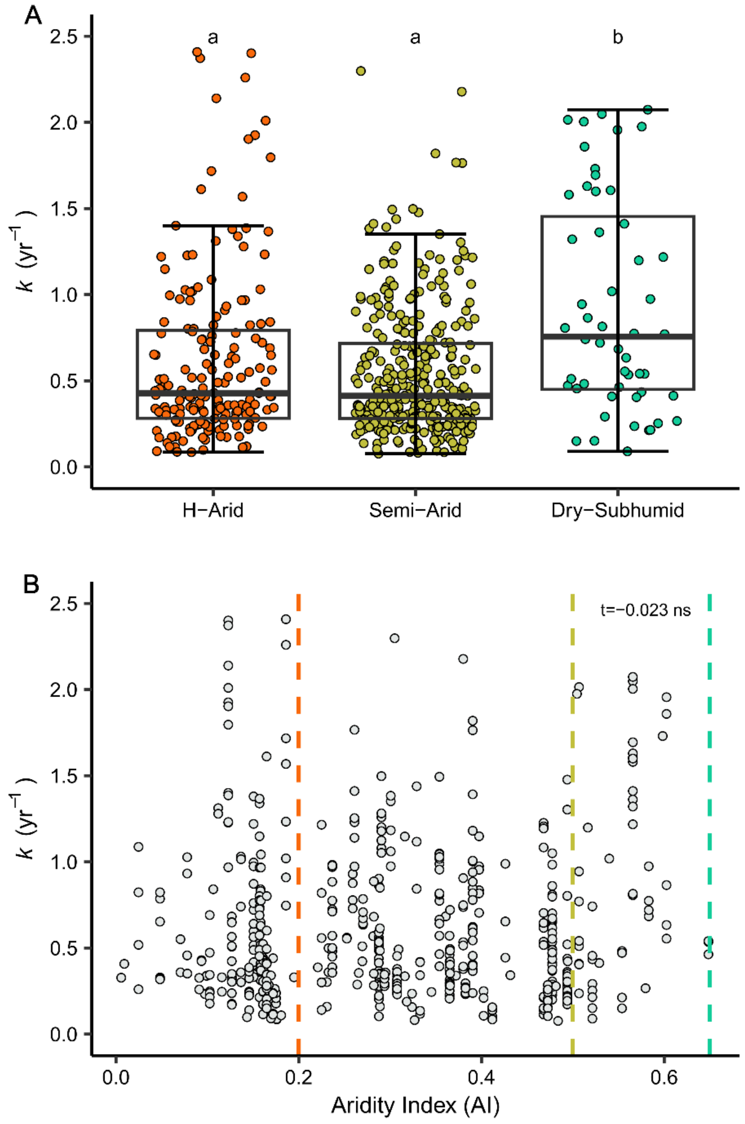

**Fig. S4. Litter decomposition correlates positively with lignin and nitrogen (N) content but does not correlate with L:N ratio globally in drylands.** (A) Lignin correlation was significant and positive for global drylands and for H-Arid and Semi-Arid classes. (B) N correlation was significant and positive for global and the three AI classes. (C) As opposite with the observed separately for Lignin and N, decomposition rates are not associated with L:N ratio for global drylands, and neither for H-Arid and Semi-Arid drylands. Conversely, Dry-Subhumid drylands showed a negative relationship. Kendall's correlation coefficients (t are presented for global data and for H-Arid, Semiarid and Dry-Subhumid categories.

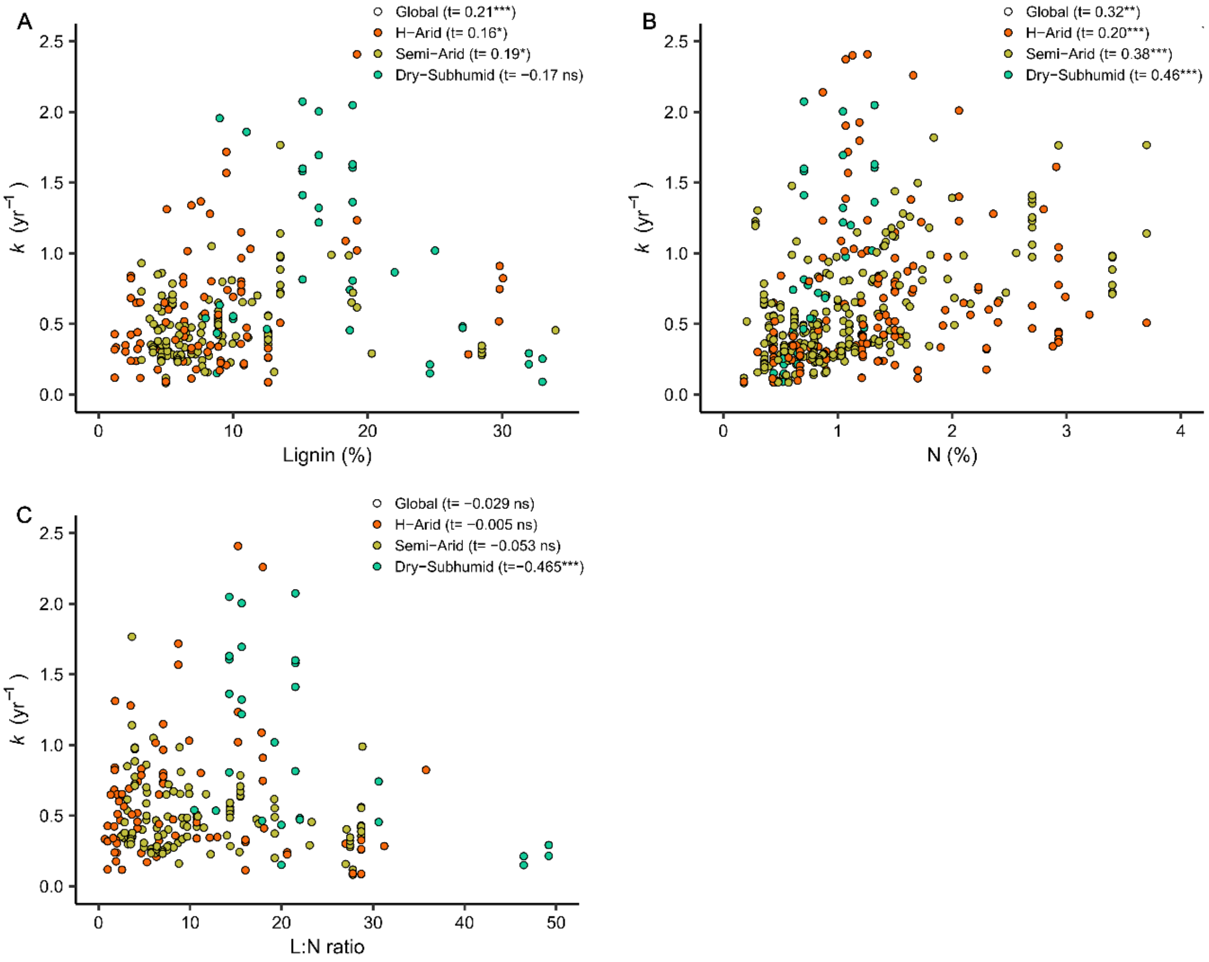

**Fig. S5. Correlations of residuals of global climatic model with litter quality.** (A) Nitrogen (%), (B) Lignin (%). Each circle represents residual values for each observation. Kendall

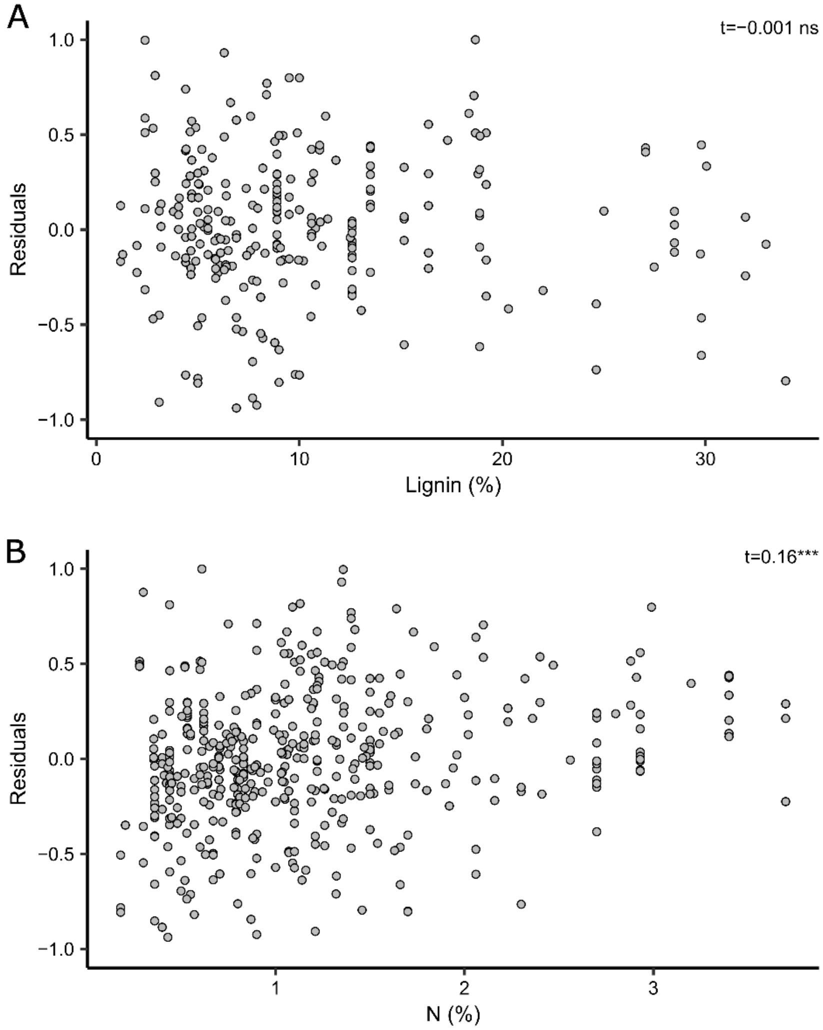

**Fig. S6. Geographical patterns of the four variables included in the global model just for drylands.** (A) MAT; (B) SEASON; (C) CVp; (D) CLOUDC.

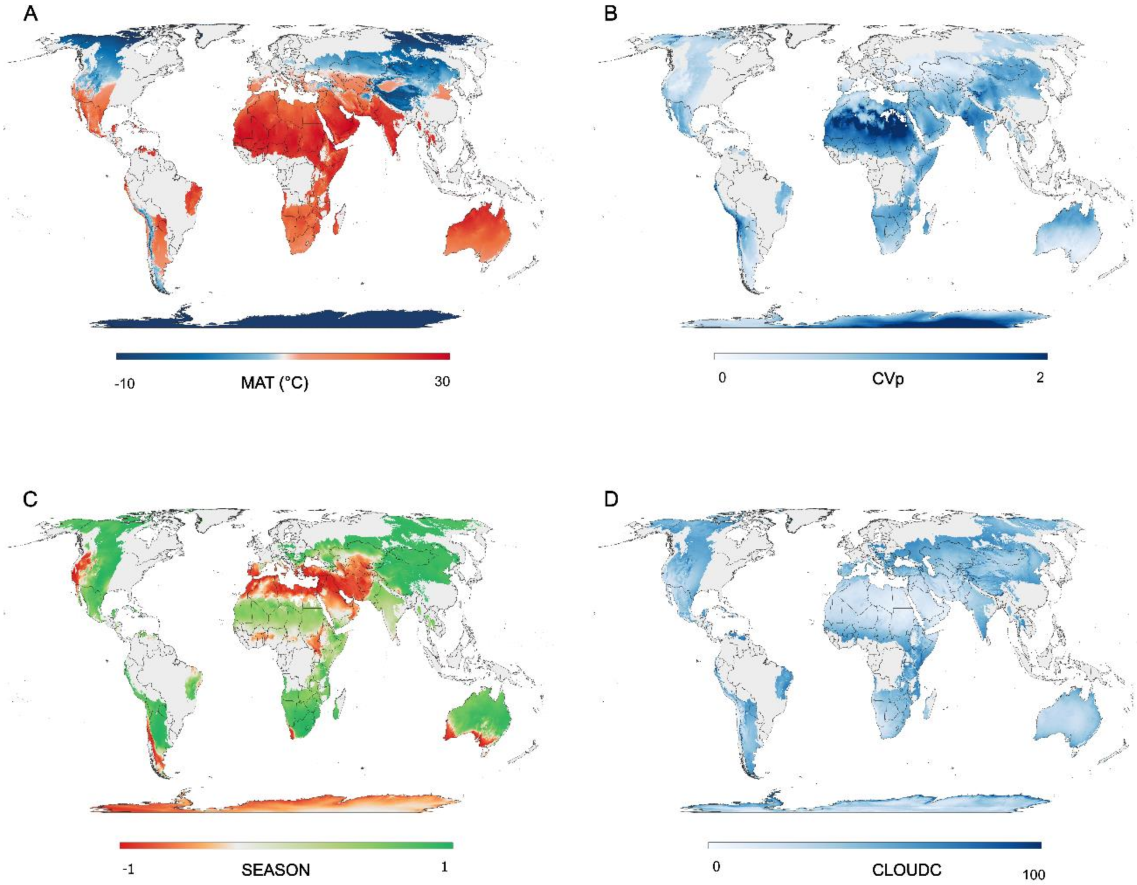

**Fig. S7. Random sampling of predicted decomposition rates across global drylands.** Map showing 12,000 randomly sampled points used to extract predicted decomposition rates ($k$, $yr^{-1}$) and associated environmental variables, including Aridity Index (AI), precipitation-temperature synchrony (SEASON), and mean annual temperature (MAT). The sampling is designed to better visualize the predicted patterns for arid regions globally, based on model outputs.

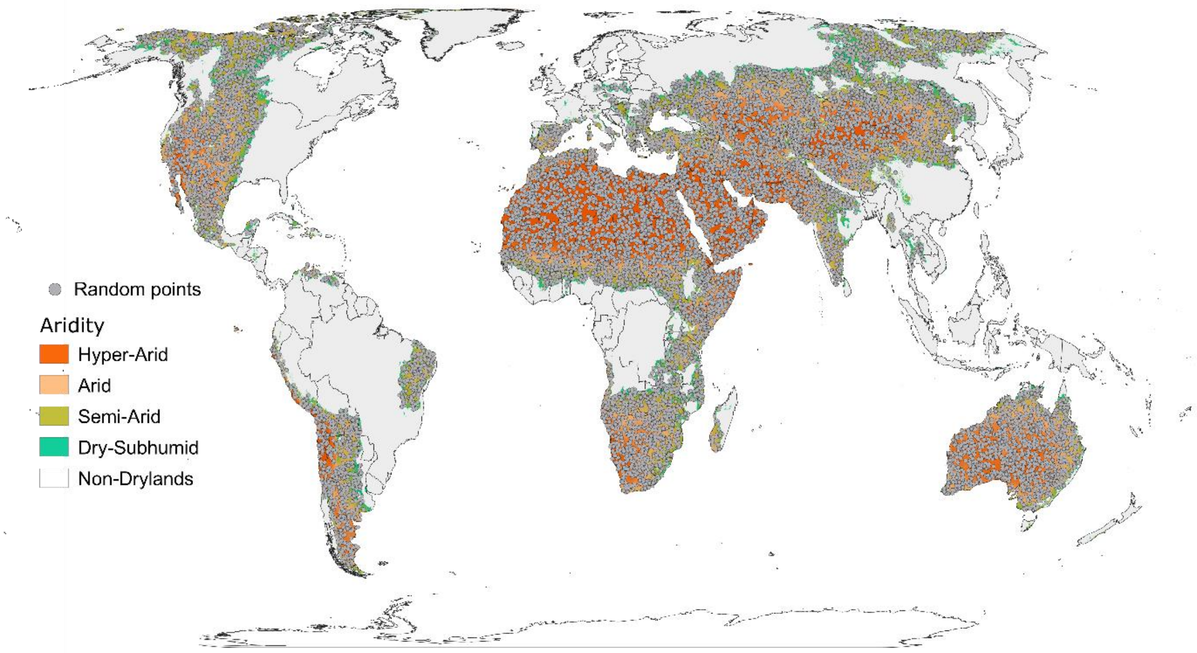

**Table S1 - Dryland sites included in analyses.** Data shown are site name (SITE), references (ID), location (COUNTRY, LATITUDE, LONGITUDE), Aridity index (AI), aridity class and number of *k* values (n).

| SITE | ID | COUNTRY | LATITUDE | LONGITUDE | AI | Aridity class | n (*k* values) |
|---|---|---|---|---|---|---|---|
| **Achala** | (1, 2) | Argentina | -31.57 | -64.83 | 0.48 | Semi-arid | 36 |
| **Afgoi** | (3) | Somalia | 2.1666667 | 45.0833333 | 0.30 | Semi-arid | 2 |
| **Aranjuez** | (4, 5) | Spain | 40.0166667 | -3.51666667 | 0.33 | Semi-arid | 5 |
| **Broken Hill** | (6) | Australia | -31.072 | 141.799 | 0.12 | H-Arid | 2 |
| **BSW** | (7, 8) | China | 44.3666667 | 87.9166667 | 0.16 | H-Arid | 6 |
| **Bulawayo** | (9) | Zimbabwe | -20.133 | 28.6 | 0.37 | Semi-arid | 3 |
| **Carlsbad** | (10) | USA | 32.408 | -104.23 | 0.18 | H-Arid | 3 |
| **Central Parkland** | (11) | Canada | 53 | -111 | 0.40 | Semi-arid | 6 |
| **Central Plains** | (12) | USA | 40.8 | -104.8 | 0.32 | Semi-arid | 2 |
| **Central Plains2** | (13) | USA | 40.82 | -104.77 | 0.32 | Semi-arid | 1 |
| **Chaco serrano** | (14) | Argentina | -31.1666667 | -64 | 0.58 | Subhumid | 1 |
| **Chancani** | (15, 16) | Argentina | -31.4 | -65.55 | 0.33 | Semi-arid | 3 |
| **Claremont** | (17) | USA | 34.1 | -117.7 | 0.26 | Semi-arid | 4 |
| **Cobar** | (18) | Australia | -32.583 | 145.586 | 0.18 | Semi-arid | 1 |
| **Córdoba** | (19) | Argentina | -32.8333333 | -63.7333333 | 0.60 | Subhumid | 1 |
| **Deerlodge Park** | (20) | USA | 40.45 | -108.52 | 0.27 | Semi-arid | 5 |
| **Duolun** | (21) | China | 42.03 | 116.283 | 0.47 | Semi-arid | 6 |

| | | | | | | | |
|---|---|---|---|---|---|---|---|
| **El Carrizal** | (22) | Mexico | 24.13 | -110.43 | 0.12 | H-Arid | 20 |
| **Givat** | (23) | Israel | 31.7845778 | 35.0888028 | 0.27 | Semi-arid | 2 |
| **Glassman20181** | (24) | USA | 33.648 | -116.38 | 0.10 | H-Arid | 1 |
| **Glassman20182** | (24) | USA | 33.61 | -116.45 | 0.19 | H-Arid | 1 |
| **Glassman20183** | (24) | USA | 33.737 | -117.7 | 0.28 | Semi-arid | 1 |
| **Glassman20184** | (24) | USA | 33.683 | -116.77 | 0.26 | Semi-arid | 1 |
| **Glassman20185** | (24) | USA | 33.823 | -116.75 | 0.48 | Semi-arid | 1 |
| **Gonglushan** | (25) | China | 36.4233333 | 109.5255 | 0.55 | Subhumid | 4 |
| **Great Basin** | (26) | USA | 39.85 | -119.38 | 0.14 | H-Arid | 3 |
| **Gruya** | (27) | Palestine | 31.5918444 | 35.4058639 | 0.09 | H-Arid | 1 |
| **Gurbantunggut** | (28) | China | 44.378 | 87.928 | 0.15 | H-Arid | 17 |
| **Har-Gilo** | (27) | Palestine | 31.7206583 | 35.1747611 | 0.27 | Semi-arid | 1 |
| **Hermosillo** | (29) | Mexico | 29.017 | -110.95 | 0.19 | H-Arid | 9 |
| **Hidalgo** | (30) | Mexico | 20.2333333 | -98.3833333 | 0.52 | Subhumid | 2 |
| **Horqin** | (31) | China | 42.6666667 | 120.933333 | 0.48 | Semi-arid | 1 |
| **Horqin2** | (32) | China | 42.917 | 120.683 | 0.43 | Semi-arid | 3 |
| **Horqin3** | (33, 34) | China | 42.9666667 | 122.35 | 0.49 | Semi-arid | 8 |
| **Horqin4** | (35) | China | 42.9666667 | 122.35 | 0.49 | Semi-arid | 24 |
| **Inner Mongolia2** | (36) | China | 42.0333333 | 116.266667 | 0.47 | Semi-arid | 9 |
| **Inner Mongolia3** | (37) | China | 43.2166667 | 116.233333 | 0.38 | Semi-arid | 5 |

| | | | | | | | |
|---|---|---|---|---|---|---|---|
| **Inner Mongolia4** | (38) | China | 43.5527778 | 116.674722 | 0.37 | Semi-arid | 1 |
| **Inner Mongolia5** | (39) | China | 43.6333333 | 116.633333 | 0.36 | Semi-arid | 1 |
| **InnerMongolia6** | (39) | China | 43.7333333 | 116.283333 | 0.31 | Semi-arid | 1 |
| **Irvine** | (40) | USA | 33.65 | -117.85 | 0.23 | Semi-arid | 2 |
| **Jaiba** | (41) | Brazil | -15.1508333 | -43.8238889 | 0.57 | Subhumid | 15 |
| **Jornada1** | (13, 42) | USA | 32.5 | -106.75 | 0.16 | H-Arid | 11 |
| **Jornada2** | (43) | USA | 32.51 | -106.79 | 0.16 | H-Arid | 4 |
| **Jornada3** | (44) | USA | 32.5 | -106.8 | 0.16 | H-Arid | 1 |
| **Jornada4** | (45) | USA | 32.6188889 | -106.787778 | 0.16 | H-Arid | 2 |
| **Jornada5** | (45) | USA | 32.6113889 | -106.796111 | 0.16 | H-Arid | 2 |
| **Jornada6** | (45) | USA | 32.6061111 | -106.850556 | 0.16 | H-Arid | 2 |
| **Jornada7** | (46) | USA | 32.563 | -106.76 | 0.16 | H-Arid | 3 |
| **Jornada8** | (43) | USA | 32.5994444 | -106.832222 | 0.16 | H-Arid | 8 |
| **Kalia** | (23) | Palestine | 31.715 | 35.4249639 | 0.10 | H-Arid | 2 |
| **Karamay** | (47) | China | 45.608 | 84.834 | 0.11 | H-Arid | 1 |
| **Kdoshim** | (48) | Israel | 31.8 | 35.0333 | 0.29 | Semi-arid | 20 |
| **La Campana** | (49) | Chile | -32.95 | -71.06 | 0.30 | Semi-arid | 1 |
| **La Jolla** | (50) | USA | 32.89 | -117.23 | 0.23 | Semi-arid | 2 |

| | | | | | | | |
|---|---|---|---|---|---|---|---|
| **La Pampa** | (51, 52) | Argentina | -38.75 | -63.75 | 0.29 | Semi-arid | 7 |
| **Las Lomitas** | (49) | Chile | -26.01 | -70.61 | 0.01 | H-Arid | 1 |
| **Los medanos** | (53) | USA | 32.3083333 | -103.821944 | 0.17 | H-Arid | 1 |
| **Madurai** | (54) | India | 9.917 | 78.167 | 0.52 | Subhumid | 1 |
| **Matobo** | (55, 56) | Zimbabwe | -20.383 | 28.467 | 0.35 | Semi-arid | 15 |
| **Mazandaran** | (57) | Iran | 36.3333333 | 51.75 | 0.29 | Semi-arid | 8 |
| **Metzukei Dragot** | (27) | Palestine | 31.5947944 | 35.38825 | 0.09 | H-Arid | 1 |
| **Mishor Adumim** | (23) | Palestine | 31.7987861 | 35.3372083 | 0.16 | H-Arid | 2 |
| **Mixed Prairie** | (11) | Canada | 51 | -112 | 0.58 | Subhumid | 4 |
| **Mojave** | (58) | USA | 36.8166667 | -114.75 | 0.07 | H-Arid | 2 |
| **Mopani** | (59) | South Africa | -22.417 | 30.833 | 0.29 | Semi-arid | 8 |
| **Mostoles** | (60) | Spain | 40.335667 | -3.876933 | 0.33 | Semi-arid | 1 |
| **Mpala** | (61) | Kenya | 0.2833333 | 36.8833333 | 0.54 | Subhumid | 1 |
| **Mu Us** | (62) | China | 37.7086111 | 107.226944 | 0.32 | Semi-arid | 2 |
| **Mu Us2** | (63, 64) | China | 39.494 | 110.192 | 0.39 | Semi-arid | 14 |
| **Naaka Junction** | (27) | Palestine | 31.5625 | 35.2859333 | 0.17 | H-Arid | 1 |
| **Negev** | (65) | Israel | 30.8166667 | 34.7333333 | 0.10 | H-Arid | 7 |
| **Negev2** | (66, 67) | Israel | 30.8666667 | 34.7666667 | 0.09 | H-Arid | 2 |
| **Negev3** | (66, 67) | Palestine | 31.3833333 | 34.9 | 0.18 | H-Arid | 2 |
| **Negev4** | (66, 67) | Israel | 31.7 | 35.05 | 0.25 | Semi-arid | 2 |
| **Negev5** | (66, 67) | Israel | 33 | 35.25 | 0.53 | Semi-arid | 1 |
| **Negev6** | (66, 67) | Israel | 33 | 35.2333333 | 0.49 | Semi-arid | 1 |

| | | | | | | | |
|---|---|---|---|---|---|---|---|
| **New Mexico** | (68) | USA | 32.5794444 | -106.5275 | 0.23 | Semi-arid | 6 |
| **OHP** | (69) | France | 43.97 | 5.88 | 0.65 | Subhumid | 3 |
| **Ordos** | (70) | China | 39.49 | 110.19 | 0.39 | Semi-arid | 8 |
| **Pampa Blanca** | (49) | Chile | -25.95 | -70.61 | 0.01 | H-Arid | 1 |
| **Patagonia1** | (71) | Argentina | -40.25 | -70.8 | 0.52 | Subhumid | 8 |
| **Phoenix** | (72, 73) | USA | 33.5 | -111.8 | 0.13 | H-Arid | 15 |
| **Phoenix2** | (74) | USA | 33.577 | -112.081 | 0.11 | H-Arid | 2 |
| **Pilliga** | (75) | Australia | -30.6565417 | 149.169661 | 0.40 | Semi-arid | 3 |
| **Pretoriuskop** | (59) | South Africa | -25.166 | 31.267 | 0.51 | Subhumid | 8 |
| **Quebrada de Talca** | (49) | Chile | -30.05 | -71.1 | 0.08 | H-Arid | 1 |
| **Queensland** | (76) | Australia | -24.3833333 | 148.25 | 0.37 | Semi-arid | 12 |
| **Rajkot** | (77) | India | 20.967 | 70.333 | 0.50 | Subhumid | 3 |
| **Ramat Hanadiv NP** | (78, 79) | Israel | 32.552 | 34.945 | 0.37 | Semi-arid | 7 |
| **Rio Mayo** | (80, 81) | Argentina | -45.683333 | -70.266667 | 0.15 | H-Arid | 7 |
| **Rock Valley** | (82) | USA | 36.65 | -116.183333 | 0.08 | H-Arid | 3 |
| **Santa Rita1** | (83) | USA | 31.783 | -110.844 | 0.30 | Semi-arid | 5 |
| **Santa Rita2** | (84) | USA | 31.793 | -110.885 | 0.24 | Semi-arid | 13 |
| **Santa Rita3** | (83) | USA | 31.803 | -110.863 | 0.26 | Semi-arid | 7 |
| **Satara** | (59) | South Africa | -24.392 | 31.783 | 0.38 | Semi-arid | 8 |

| | | | | | | | |
|---|---|---|---|---|---|---|---|
| **Sedgwick Reserve** | (85) | USA | 34.7 | -120.03 | 0.35 | Semi-arid | 1 |
| **Sevilleta1** | (13, 86) | USA | 34.33 | -106.67 | 0.17 | H-Arid | 6 |
| **Sevilleta2** | (12) | USA | 34.4 | -106.9 | 0.14 | H-Arid | 2 |
| **SF** | (7, 8) | China | 45.25 | 87.6 | 0.05 | H-Arid | 6 |
| **Snake River Plain** | (87) | USA | 43 | -112 | 0.43 | Semi-arid | 1 |
| **Sobral** | (88) | Brazil | -3.7 | -40.35 | 0.60 | Subhumid | 6 |
| **Songnen** | (89) | China | 44.6666667 | 123.733333 | 0.47 | Semi-arid | 8 |
| **Sorbas** | (4) | Spain | 37.0833333 | -2.06666667 | 0.17 | H-Arid | 2 |
| **South Africa** | (90) | South Africa | -24.78 | 26.17 | 0.28 | Semi-arid | 4 |
| **Steens Mountain** | (91) | USA | 42.93 | -118.61 | 0.18 | H-Arid | 1 |
| **Sunset Crater NM** | (92) | USA | 35.37 | -111.55 | 0.41 | Semi-arid | 8 |
| **Taklimakan** | (93) | China | 39.0166667 | 83.6 | 0.02 | H-Arid | 4 |
| **Teqoa** | (27) | Palestine | 31.6516528 | 35.227125 | 0.22 | Semi-arid | 1 |
| **Tianshan** | (94) | China | 42 | 83 | 0.17 | H-Arid | 1 |
| **Tianshan2** | (95) | China | 43.4936111 | 87.3019444 | 0.29 | Semi-arid | 4 |
| **Turpan** | (7, 8) | China | 42.860277 | 89.1930556 | 0.17 | H-Arid | 6 |
| **Urat Houqi** | (31) | China | 41.4166667 | 106.966667 | 0.14 | H-Arid | 1 |
| **Xilin** | (96) | China | 43.6333333 | 116.7 | 0.37 | Semi-arid | 3 |
| **Xilinhot** | (97) | China | 44.1673333 | 116.482444 | 0.31 | Semi-arid | 28 |
| **Yatir** | (48) | Israel | 31.347 | 35.0356 | 0.16 | H-Arid | 12 |

**Table S2.** Detailed description of variables used for the analyses.

| Variable | Units | Description | Provider | Database (product) | Resolution | Time period |
|---|---|---|---|---|---|---|
| MAT | °C | Mean annual temperature | TerraClimate | Climatologies | 4 km | 1981-2010 |
| MAP | Mm | Mean annual precipitation | TerraClimate | Climatologies | 4 km | 1981-2010 |
| RAD | W/m2.yr | Total accumulated solar radiation by year | TerraClimate | Climatologies | 4 km | 1981-2010 |
| SEASON | Unitless | The overlap between wet and warm seasons, which was estimated by the Pearson correlation coefficient on monthly data of precipitation and temperature. Values between -1 (mediterranean) and +1 (monsoon). | Derived from TerraClimate | Derived from monthly data of Climatologies | 4 km | 1981-2010 |
| CVp | Unitless | Precipitation variability estimated throughout coefficient of variation of interannual precipitation | Derived from Terraclimate | Derived from monthly data of Climatologies | 4 km | 1981-2010 |
| NDVI | Unitless | Mean annual normalized difference vegetation index | LPDAAC | MODIS (VNP13A3 v001) | 500 m | 2000-2020 |
| CLOUDC | Unitless | Frequency of days with cloud cover | EarthEnv | Global-1km-CloudCover | 1 km | 2000-2014 |

**Table S3. Summary of decomposition rates (*k*) and data distribution across aridity and seasonality classes.** Median and range of values of *k* were presented.

| **Aridity (IA)** | **N / site** | ***k* ($yr^{-1}$)** |
|---|---|---|
| H-Arid | 188 / 45 | 0.437a<br>(0.085-5.101) |
| Semi-Arid | 342 / 57 | 0.415a<br>(0.077-4.745) |
| Dry-Subhumid | 58 / 14 | 0.811b<br>(0.090-5.903) |
| **Seasonality** | **N / site** | ***k* ($yr^{-1}$)** |
| Mediterranean | 134 / 45 | 0.395a<br>(0.082-3.086) |
| Monsoon | 425 / 67 | 0.517b<br>(0.077-5.903) |
| Iso (Non Seasonal) | 29 / 5 | 0.328a<br>(0.085-1.199) |

**Table S4. Summary of decomposition rates (*k*), data distribution and climatic variables across ecosystems.** Median and range of values of *k* were presented. For climate variables mean and range of values are presented.

| ECOSYSTEM | n/site | *k* ($year^{-1}$) | Aridity index (AI) | MAP (mm) | MAT (°C) | CVp |
|---|---|---|---|---|---|---|
| *Desert* | 161/38 | 0.55a (0.10-4.75) | 0.16 (0.005-0.39) | 239.6 (8.7-510.2) | 15.45 (1.60-23.54) | 0.79 (0.37-1.77) |
| *Shrubland* | 57/15 | 0.72a (0.09-5.10) | 0.25 (0.08-0.60) | 388.2 (91.6-947.7) | 20.03 (13.60-27.11) | 1.18 (0.27-1.40) |
| *Grassland* | 97/22 | 0.55ab (0.08-5.80) | 0.43 (0.12-0.60) | 418.3 (220.9-945.2) | 9.66 (1.33-29.06) | 0.69 (0.24-1.70) |
| *Savanna* | 62/10 | 0.51ab (0.10-3.03) | 0.36 (0.18-0.54) | 561.0 (342.6-749.5) | 21.13 (18.08-27.18) | 0.84 (0.21-1.02) |
| *Forest* | 134/23 | 0.35bc (0.09-5.90) | 0.41 (0.10-0.65) | 466.6 (127.6-803.5) | 9.90 (4.10-24.50) | 1.02 (0.26-1.23) |
| *Steppe* | 77/12 | 0.30c (0.10-1.23) | 0.37 (0.15-0.52) | 307.4 (188.2-630.5) | 3.48 (1.55-9.96) | 1.14 (0.21-1.24) |

**Tabla S5. Top 30 candidate models explaining the relationship between ln *k* (decomposition rate) and climate variables.** Models are ranked by ΔAIC relative to the global model (Model 1). The table includes parameters, coefficients of fixed effects and intercept, degrees of freedom (df), AICc, log-likelihood, and model weight.

| Nm | Intercept | MAT | SEASON | CVp | CLOUDC | MAP | NDVI | RAD | df | logLik | AICc | delta | Weight |
|---|---|---|---|---|---|---|---|---|---|---|---|---|---|
| **1** | -0.75 | 0.35 | 0.15 | 0.11 | 0.15 | | | | 7 | -451.53 | 917.26 | 0.00 | 0.13 |
| **2** | -0.75 | 0.37 | 0.15 | 0.11 | 0.18 | | -0.06 | | 8 | -451.14 | 918.52 | 1.26 | 0.07 |
| **3** | -0.76 | 0.35 | 0.14 | | 0.13 | | | | 6 | -453.24 | 918.62 | 1.36 | 0.07 |
| **4** | -0.75 | 0.33 | 0.13 | 0.11 | 0.13 | 0.14 | -0.15 | | 9 | -450.26 | 918.82 | 1.56 | 0.06 |
| **5** | -0.76 | 0.26 | 0.14 | 0.10 | | 0.22 | -0.15 | | 8 | -451.48 | 919.20 | 1.94 | 0.05 |
| **6** | -0.75 | 0.33 | 0.15 | 0.11 | 0.13 | 0.02 | | | 8 | -451.48 | 919.21 | 1.95 | 0.05 |
| **7** | -0.75 | 0.34 | 0.15 | 0.11 | 0.15 | | | 0.00 | 8 | -451.53 | 919.31 | 2.06 | 0.05 |
| **8** | -0.75 | 0.26 | 0.16 | 0.10 | | 0.10 | | | 7 | -452.73 | 919.64 | 2.39 | 0.04 |
| **9** | -0.77 | 0.27 | 0.14 | | | 0.21 | -0.15 | | 7 | -452.87 | 919.92 | 2.67 | 0.04 |
| **10** | -0.76 | 0.38 | 0.14 | | 0.16 | | -0.05 | | 7 | -452.93 | 920.06 | 2.80 | 0.03 |
| **11** | -0.76 | 0.30 | 0.18 | 0.09 | | | | | 6 | -454.00 | 920.14 | 2.88 | 0.03 |
| **12** | -0.76 | 0.27 | 0.15 | | | 0.10 | | | 6 | -454.04 | 920.22 | 2.96 | 0.03 |
| **13** | -0.76 | 0.33 | 0.12 | | 0.11 | 0.15 | -0.14 | | 8 | -452.01 | 920.27 | 3.01 | 0.03 |
| **14** | -0.76 | 0.34 | 0.15 | 0.10 | | 0.21 | -0.16 | -0.09 | 9 | -451.07 | 920.44 | 3.19 | 0.03 |
| **15** | -0.76 | 0.33 | 0.14 | | 0.11 | 0.03 | | | 7 | -453.15 | 920.49 | 3.24 | 0.03 |
| **16** | -0.77 | 0.31 | 0.17 | | | | | | 5 | -455.20 | 920.51 | 3.25 | 0.03 |
| **17** | -0.75 | 0.38 | 0.15 | 0.11 | 0.18 | | -0.06 | 0.00 | 9 | -451.14 | 920.58 | 3.33 | 0.03 |
| **18** | -0.76 | 0.34 | 0.14 | | 0.14 | | | 0.02 | 7 | -453.22 | 920.64 | 3.38 | 0.02 |
| **19** | -0.75 | 0.35 | 0.13 | 0.11 | 0.12 | 0.15 | -0.15 | -0.02 | 10 | -450.23 | 920.84 | 3.59 | 0.02 |
| **20** | -0.76 | 0.39 | 0.18 | 0.10 | | | | -0.11 | 7 | -453.37 | 920.94 | 3.68 | 0.02 |
| **21** | -0.75 | 0.33 | 0.16 | 0.10 | | 0.09 | | -0.07 | 8 | -452.47 | 921.18 | 3.92 | 0.02 |
| **22** | -0.75 | 0.33 | 0.15 | 0.11 | 0.13 | 0.02 | | 0.00 | 9 | -451.48 | 921.27 | 4.02 | 0.02 |
| **23** | -0.78 | 0.28 | | 0.10 | 0.15 | 0.19 | -0.19 | | 8 | -452.60 | 921.44 | 4.18 | 0.02 |
| **24** | -0.78 | 0.30 | | 0.10 | 0.19 | | | | 6 | -454.67 | 921.49 | 4.23 | 0.02 |
| **25** | -0.77 | 0.33 | 0.14 | | | 0.20 | -0.15 | -0.07 | 8 | -452.65 | 921.54 | 4.28 | 0.02 |
| **26** | -0.77 | 0.38 | 0.17 | | | | | -0.09 | 6 | -454.81 | 921.77 | 4.51 | 0.01 |
| **27** | -0.76 | 0.32 | 0.15 | | | 0.09 | | -0.05 | 7 | -453.92 | 922.03 | 4.77 | 0.01 |
| **28** | -0.76 | 0.37 | 0.14 | | 0.16 | | -0.05 | 0.01 | 8 | -452.92 | 922.10 | 4.84 | 0.01 |
| **29** | -0.76 | 0.30 | 0.18 | 0.09 | | | 0.01 | | 7 | -453.97 | 922.13 | 4.87 | 0.01 |
| **30** | -0.79 | 0.31 | | | 0.18 | | | | 5 | -456.03 | 922.15 | 4.90 | 0.01 |
| **RELATIVE IMPORTANCE (sum of weights)** | | 1 | 0.96 | 0.66 | 0.67 | 0.45 | 0.41 | 0.26 | | | | | |

**Table S6. Global model testing the relationship between decomposition rate (*k*) and climate variables.** The model is based on 588 *k* observations across 116 sites. Estimates represent the mean values of fixed effects coefficients, with 95% confidence intervals (CI). Significance levels are indicated by p-values, with *p < 0.05 and **p < 0.01. Random effects include site (n = 116). The intercept represents the mean value of *k* when all predictors are set to 0. The table presents parameter estimates, confidence intervals (CI-95%), p-values, Akaike Information Criterion (AIC), semi-partial $R^2$ (part$R^2$), inclusive $R^2$ ($R^2$Inc), and Variance Inflation Factors (VIF) for each predictor. Marginal $R^2$ ($R^2$m = 0.17) and conditional $R^2$ ($R^2$c = 0.72) are provided to indicate the explained variance by fixed and combined fixed/random effects, respectively.

**GLOBAL MODEL**

| **Predictors** | **Estimates** | **CI (95%)** | **P value** | **part$R^2$ / $R^2$Inc** | **VIF** |
|---|---|---|---|---|---|
| Intercept | -0.75 | (-0.87 – -0.62) | <0.001 | | |
| MAT | 0.35 | (0.20 – 0.49) | <0.001 | 0.136 / 0.091 | 1.23 |
| SEASON | 0.15 | (0.03 – 0.27) | 0.013 | 0.037 / 0.008 | 1.23 |
| CVp | 0.11 | (-0.01 – 0.23) | 0.071 | 0.018 / 0.018 | 1.04 |
| CLOUDC | 0.15 | (0.02 – 0.28) | 0.029 | 0.022 / 0.003 | 1.26 |
| **Sites** | 116 | | | | |
| **Obs (n)** | 588 | | | | |
| **$R^2$m** | 0.17 | | | | |
| **$R^2$c** | 0.72 | | | | |

**Table S7. Subset models testing the relationship between decomposition rate (*k*) and climate variables for different aridity classes.** Each model corresponds to observations from the three aridity classes. Estimates represent the mean values of fixed effects coefficients, with 95% confidence intervals (CI). Significance levels are indicated by p-values, with $*p < 0.05$ and $**p < 0.01$. Random effects include site (H-Arid: n = 45; Semi-Arid: n = 58; Dry-Subhumid: n = 13). The intercept represents the mean value of *k* when all predictors are set to 0. The table presents parameter estimates, confidence intervals (CI-95%), p-values, Akaike Information Criterion (AIC), semi-partial $R^2$ (part$R^2$), inclusive $R^2$ ($R^2$Inc), and Variance Inflation Factors (VIF) for each predictor. Marginal $R^2$ ($R^2$m) and conditional $R^2$ ($R^2$c) are reported to indicate the explained variance by fixed and combined fixed/random effects, respectively.

| **H-ARID MODEL** | | | | | |
|---|---|---|---|---|---|
| **Predictors** | **Estimates** | **CI (95%)** | **P value** | **$R^2$semi-partial / R2 Inclusive** | **VIF** |
| Intercept | -0.82 | (-1.00 – 0.64) | <0.001 | | |
| MAT | 0.37 | (0.13 – 0.62) | 0.003 | 0.170 / 0.085 | 1.37 |
| SEASON | 0.27 | (0.10 – 0.45) | 0.003 | 0.143 / 0.036 | 1.28 |
| MAP | -0.14 | (-0.30 – 0.03) | 0.108 | 0.044 / 0.001 | 1.18 |
| **Sites** | 45 | | | | |
| **Obs (n)** | 188 | | | | |
| **$R^2$m** | 0.2 | | | | |
| **$R^2$c** | 0.63 | | | | |

| **SEMI-ARID MODEL** | | | | | |
|---|---|---|---|---|---|
| **Predictors** | **Estimates** | **CI (95%)** | **P value** | **$R^2$ semi-partial / R2 Inclusive** | **VIF** |
| Intercept | -0.85 | (-1.02 – 0.68) | <0.001 | | |
| MAT | 0.39 | (0.18 – 0.60) | 0.001 | 0.100 / 0.043 | 1.70 |
| CVp | 0.14 | (-0.02 – 0.30) | 0.075 | 0.019 / 0.008 | 1.12 |
| CLOUDC | 0.16 | (-0.03 – 0.34) | 0.091 | <0.0001 / 0.005 | 1.33 |
| NDVI | -0.24 | (-0.42 – -0.07) | 0.006 | 0.041 / 0.002 | 1.44 |
| **Sites** | 57 | | | | |
| **Obs (n)** | 342 | | | | |
| **$R^2$m** | 0.13 | | | | |

| | | | | | |
|---|---|---|---|---|---|
| **$R^2$c** | 0.71 | | | | |
| | | **DRY-SUBHUMID MODEL** | | | |
| **Predictors** | **Estimates** | **CI (95%)** | **P value** | **R2 semi-partial / R2 Inclusive** | **VIF** |
| Intercept | -0.21 | (-0.64 – -0.21) | <0.001 | - | - |
| RAD | 0.49 | (0.03 - 0.88) | 0.039 | - | - |
| **Sites** | 14 | | | | |
| **Obs (n)** | 58 | | | | |
| **$R^2$m** | 0.21 | | | | |
| **$R^2$c** | 0.74 | | | | |

**Table S8. Lignin Model testing the relationship between decomposition rate (*k*) and lignin content.** Estimates represent the mean values of fixed effects coefficients, with 95% confidence intervals (CI). Significance levels are indicated by p-values. The intercept represents the mean *k* value when all predictors are 0. The model includes species as a random effect (n = 111 species; 254 observations). Marginal $R^2$ ($R^2m = 0.101$) and conditional $R^2$ ($R^2c = 0.553$) indicate the variance explained by fixed and combined fixed/random effects, respectively. ln*k* ($yr^{-1}$) = LIGNIN (%) + LIGNIN_broken + (1|SPECIES).

| **LIGNIN MODEL** | | | |
|---|---|---|---|
| **Predictors** | **Estimates** | **CI (95%)** | **P value** |
| Intercept | -1.15 | (-1.36 – -0.93) | <0.001 |
| LIGNIN | 0.05 | (0.03 – 0.07) | <0.001 |
| LIGNIN BROKEN | -0.10 | (-0.15 – -0.05) | <0.001 |
| **Species** | 111 | | |
| **Obs (n)** | 254 | | |
| **$R^2$m** | 0.101 | | |
| **$R^2$c** | 0.553 | | |

**Table S9.** Estimates represent the mean values of fixed effects coefficients, with 95% confidence intervals (CI). Significance levels are indicated by p-values. The intercept represents the mean *k* value when all predictors are 0. The model includes species as a random effect (n = 180 species; 452 observations). Marginal $R^2$ ($R^2m$ = 0.096) and conditional $R^2$ ($R^2c$ = 0.625) indicate the variance explained by fixed and combined fixed/random effects, respectively. ln*k* ($yr^{-1}$) = NITROGEN (%) + (1|SPECIES).

| **NITROGEN MODEL** | | | |
|---|---|---|---|
| **Predictors** | **Estimates** | **CI (95%)** | **P value** |
| Intercept | -0.97 | (-1.15 – -0.79) | **<0.001** |
| N | 0.30 | (0.18 – 0.43) | **<0.001** |
| **Species** | 180 | | |
| **Obs (n)** | 452 | | |
| **$R^2$m** | 0.096 | | |
| **$R^2$c** | 0.625 | | |
